# MTCL2 promotes asymmetric microtubule organization by crosslinking microtubules on the Golgi membrane

**DOI:** 10.1101/2021.09.07.459224

**Authors:** Risa Matsuoka, Masateru Miki, Sonoko Mizuno, Yurina Ito, Chihiro Yamada, Atsushi Suzuki

## Abstract

The Golgi complex plays an active role in organizing asymmetric microtubule arrays essential for polarized vesicle transport. The coiled-coil protein MTCL1 stabilizes microtubules nucleated from the Golgi membrane. Here, we report an MTCL1 paralog, MTCL2, which preferentially acts on the perinuclear microtubules accumulated around the Golgi. MTCL2 associates with the Golgi membrane through the N-terminal coiled-coil region and directly binds microtubules through the conserved C-terminal domain without promoting microtubule stabilization. Knockdown of MTCL2 significantly impaired microtubule accumulation around the Golgi as well as the compactness of the Golgi ribbon assembly structure. Given that MTCL2 forms parallel oligomers through homo-interaction of the central coiled-coil motifs, our results indicate that MTCL2 promotes asymmetric microtubule organization by crosslinking microtubules on the Golgi membrane. Results of *in vitro* wound healing assays further suggest that this function of MTCL2 enables integration of the centrosomal and Golgi-associated microtubules on the Golgi membrane, supporting directional migration. Additionally, the results demonstrated the involvement of CLASPs and giantin in mediating the Golgi association of MTCL2.

## Introduction

The microtubule (MT) cytoskeleton plays an essential role in organizing intracellular structures by mediating the transport and positioning of organelles. Generally, animal cells radiate MTs from the centrosome, where MT nucleation and attachment of MT minus ends occur predominantly (Vorobjev & Nadezhdina, 1987; Conduit *et al*, 2015). However, accumulating evidence has demonstrated that cultured cells also develop non-centrosomal MTs that nucleate from or attach their minus ends to the Golgi membrane (Wu *et al*, 2016; Efimov *et al*, 2007; Rivero *et al*, 2009; Nishita *et al*, 2017; Meiring *et al*, 2020). In contrast to centrosomal MTs, which exhibit dynamic instability at their plus ends, Golgi-associated MTs are specifically stabilized (Chabin-Brion *et al*, 2001; Bartolini & Gundersen, 2006; Rivero *et al*, 2009) and connect the individual Golgi stacks laterally (Miller *et al*, 2009). This connection leads to the formation of vertebrate-specific, crescent-like assembly of Golgi stacks, called the Golgi ribbon (Miller *et al*, 2009), required for the polarization of vesicle transport and directional migration (Miller *et al*, 2009; Wei & Seemann, 2010; Yadav *et al*, 2009a)

The molecular mechanisms by which Golgi-associated MTs nucleate from or attach their minus ends to the Golgi membrane have been studied extensively. CLASPs and AKAP450 promote microtubule nucleation from the Golgi membrane, whereas CAMSAPs are involved in the attachment of MT minus ends to the Golgi membrane (Rivero *et al*, 2009; Efimov *et al*, 2007; Wu *et al*, 2016; Wu & Akhmanova, 2017; Yang *et al*, 2017; Sanders & Kaverina, 2015). Until recently, however, the specific mechanism of stabilization of MTs was not well clarified. We identified a novel MT-regulating protein named microtubule crosslinking factor 1 (MTCL1) that specifically condenses to the Golgi membrane (Sato *et al*, 2013, 2014). MTCL1 is a long coiled-coil protein with two MT-binding domains (MTBDs) at the N- and C-terminal regions (Fig. 1A), the latter of which has a unique ability to stabilize the polymerization state of MTs (Abdul Kader *et al*, 2017; Sato *et al*, 2014, 2013). By associating with the Golgi membrane through the interaction with CLASPs and AKAP450, MTCL1 plays an essential role in the stabilization of Golgi-associated MTs through this C-MTBD activity. MTCL1 is suggested to form parallel dimers via the coiled-coil-rich region and crosslinks Golgi-associated MTs through N-MTBD lacking MT stabilizing activity (Abdul Kader et al., 2017).

**Figure 1.**
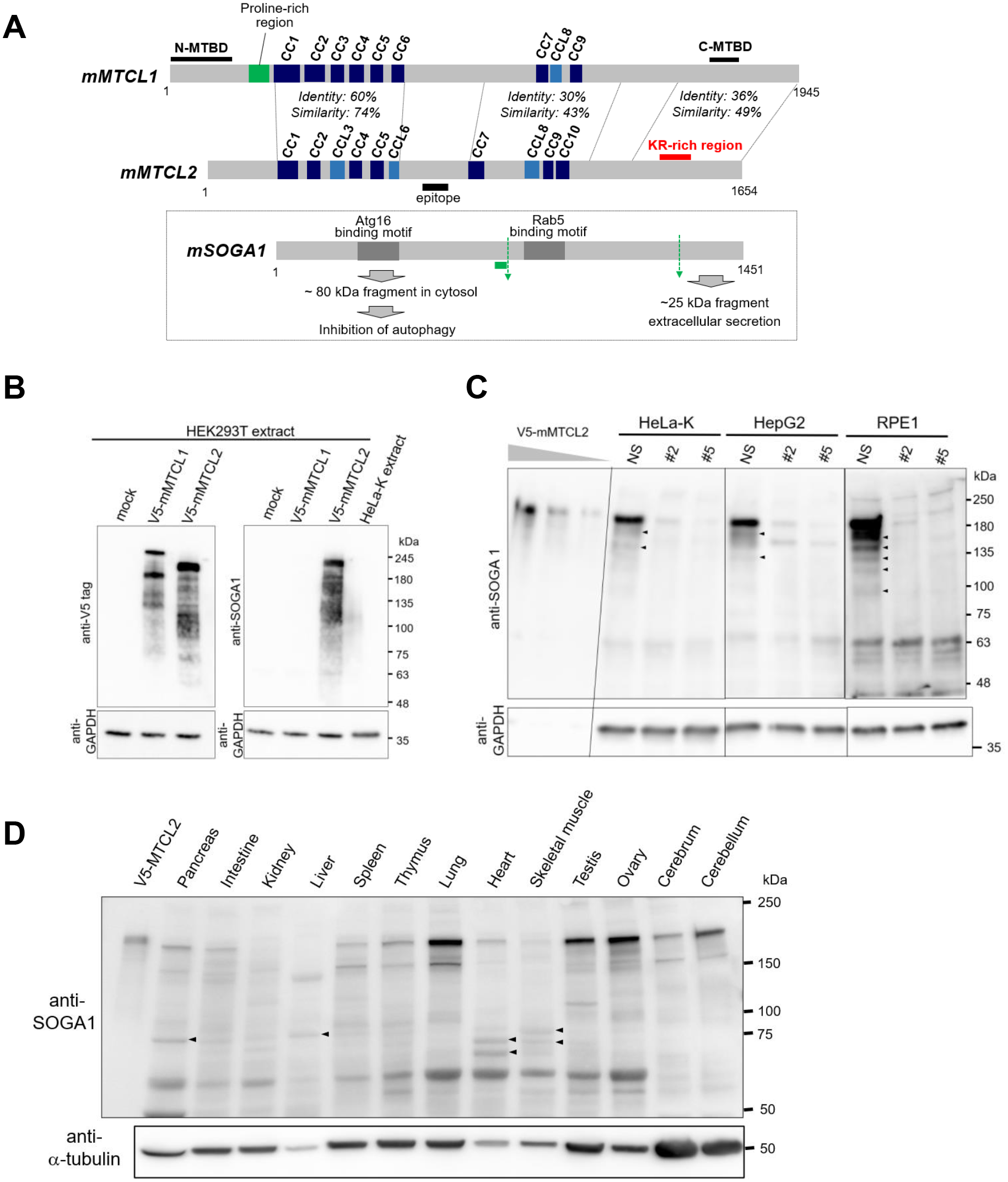
MTCL2 is expressed predominantly as a 180 kDa full-length uncleaved protein. A. Predicted molecular structure of mouse MTCL2 (mMTCL2) and its amino acid sequence homology with mouse MTCL1 (mMTCL1). CC (dark blue) corresponds to the region with the highest score (>0.85) of coiled-coil prediction, whereas CCL (light blue) corresponds to the region with a moderate score (>0.4) (https://embnet.vital-it.ch/software/COILS_form.html). The black bar labeled “epitope” indicates the position of the antigen peptide of the anti-SOGA1 antibody used in this study. The red bar indicates the region named KR-rich region whose amino acid sequence shows significant homology with C-MTBD of MTCL1 (Fig. EV1C). The boxed illustrations at the bottom indicate the structure of mouse SOGA1 (mSOGA1) in comparison with full-length mMTCL2 and summarize the arguments presented in the study reporting SOGA1 (Cowherd *et al*, 2010). The green bar indicates the predicted position of the internal signal sequence, whereas green dotted arrows indicate the predicted positions of cleavages. B. Western blotting analysis of HEK293T extracts transfected with indicated expression vectors. Used antibodies are indicated on the left of each panel. Note that detection was performed at a low sensitivity, wherein endogenous MTCL2 could not be detected in HEK293T or HeLa-K cell extracts (see lane 1 or 4 in the right panel, respectively). C. Western blotting analysis of endogenous MTCL2 in various cultured cells using an anti-SOGA1 antibody. In lane 1–3, cell extracts of HEK293T expressing exogenous V5-mMTCL2 were loaded after serial dilutions (1/10, 1/30, and 1/100). In other lanes, extracts of indicated culture cells with or without MTCL2 knockdown were loaded. NS: non-silencing control; #2 and #5 indicate different siRNAs for MTCL2. D. Tissue distribution of MTCL2. Total extracts from the indicated mouse tissues (25 μg/lane) were loaded for western blotting analysis using an anti-SOGA1 antibody. In lane 1, total cell extracts of HEK293T expressing exogenously expressed V5-mMTCL2 were loaded as a positive control.

These MTCL1 functions are specifically utilized in vertebrates because invertebrate genomes do not encode proteins homologous to MTCL1. Contrastingly, a single paralog of MTCL1, named MTCL2, is encoded in vertebrate genomes (GenBank accession number: NM_001164663). The deduced amino acid sequence of MTCL2 showed significant homology with MTCL1 in the coiled-coil region and the C-MTBD but not in the N-MTBD (Fig. 1A, Fig. EV1). This result suggests that vertebrates exploit other MT-regulating proteins with similar, but not identical, activity to that of MTCL1. Contrary to this prediction, a shorter isoform of mouse MTCL2 lacking the 203 N-terminal amino acids has already been reported as a suppressor of glucose from autophagy (SOGA) with completely different functions from those of MTCL1 (Fig. 1A) (Cowherd *et al*, 2010; Combs & Marliss, 2014). SOGA (now called SOGA1) is translated as a membrane-spanning protein and cleaved into two halves in the ER of hepatocytes (Cowherd *et al*, 2010). The resultant N-terminal fragment is released into the cytoplasm to suppress autophagy by interacting with the Atg5/Atg12/Atg16 complex, whereas the C-terminal fragment is secreted after further cleavage (Fig. 1A).

In this study, we first analyzed the expression, subcellular localization, and functions of MTCL2 and demonstrated that uncleaved MTCL2 was expressed ubiquitously and functioned as a functional paralogue of MTCL1 in the cytosol. Structure-function analysis indicated that MTCL1 forms parallel oligomers through the central coiled-coil region and crosslinks MTs by direct interaction via the C-terminal region lacking MT-stabilizing activity. In contrast to MTCL1, the Golgi association region was distinctly confined to the N-terminal coiled-coil region, which interacted with CLASP2. The involvement of giantin in the association of MTCL2 with Golgi has also been suggested. Knockdown experiments revealed that these activities of MTCL2 were required for MT accumulation around the Golgi and the clustering of Golgi stacks into a compact Golgi ribbon. *In vitro* wound healing assays further suggested a possible function of MTCL2 in integrating the centrosomal and Golgi-associated MTs around the Golgi ribbon, thus playing essential roles in directional migration. These results indicate the important roles of MTCL2 in asymmetrically organizing MTs based on the Golgi complex.

## Results

### MTCL2 is expressed predominantly as a 180 kDa full-length uncleaved protein

A mouse MTCL2 (mMTCL2) isoform lacking the 203 N-terminal amino acids, named SOGA1, was reported to be cleaved into several fragments on the ER (Fig. 1A) (Cowherd *et al*, 2010). If this processing occurs for full-length MTCL2, it cannot serve as a functional paralog of MTCL1. Thus, we first analyzed the molecular mass of MTCL2 in cultured cells using a commercially available anti-SOGA1 antibody, predicted to detect an ∼80 kDa cleaved product derived from the MTCL2 N-terminus (Fig. 1A). Results of western blotting analysis of HEK293T cells transfected with an expression vector harboring full-length mMTCL2 cDNA with an N-terminal V5-tag sequence is shown in Fig. 1B. Under a low sensitive condition at which the anti-SOGA1 antibody revealed no bands in the lanes of untransfected cells (see lanes indicated with “mock” or “HeLa-K extract”), a single major band corresponding to a molecular mass of approximately 200 kDa (Fig. 1B, right panel) was detected in the lane of V5-MTCL2-expressing cells. This molecular mass is close to the nominal molecular weight of 185.66 kDa predicted for the full-length mMTCL2 product. A similar band was detected using an anti-V5 antibody, indicating that this band corresponds to the major product derived from the transfected cDNA (Fig. 1B, left panel). Additionally, reactions with both antibodies yielded smeared bands corresponding to the molecular weights ranging from 100 to 180 kDa; however, no clear bands around 80 kDa were detected. Taken together, we concluded that full-length MTCL2 exogenously expressed in HEK293T cells was not subjected to significant intramolecular cleavage as previously reported.

Next, we examined the molecular mass of endogenous MTCL2 in extracts from several cell lines, including a human liver cancer cell line, HepG2, using the same anti-SOGA1 antibody under higher sensitive conditions (Fig. 1C). Under these conditions, a major band corresponding to ~200 kDa was detected in the lanes of HeLa-K, HepG2, and RPE1 cells. These bands corresponded to a molecular mass similar to that of V5-mMTCL2 exogenously expressed in HEK293T cells and were not observed in the lanes of cells subjected to MTCL2 knockdown (Fig. 1C). Considering that the antibody did not cross-react with MTCL1 (Fig. 1B, right panel), these results demonstrated that the examined cell lines dominantly expressed full-length MTCL2. As shown in Fig. 1C, several minor bands corresponding to molecular masses lower than 200 kDa (arrowheads) were absent in knockdown cells, particularly in RPE1 cells (Fig. 2C), suggesting that they correspond to splicing isoforms or cleaved products of MTCL2. However, most of the bands corresponded to molecular masses greater than 100 kDa, and clear bands at ∼80 kDa corresponding to the N-terminal cleavage product were not detected. Collectively, these results indicate that endogenous MTCL2 in these cell lines was not subjected to cleavage, as previously reported for SOGA1.

**Figure 2.**
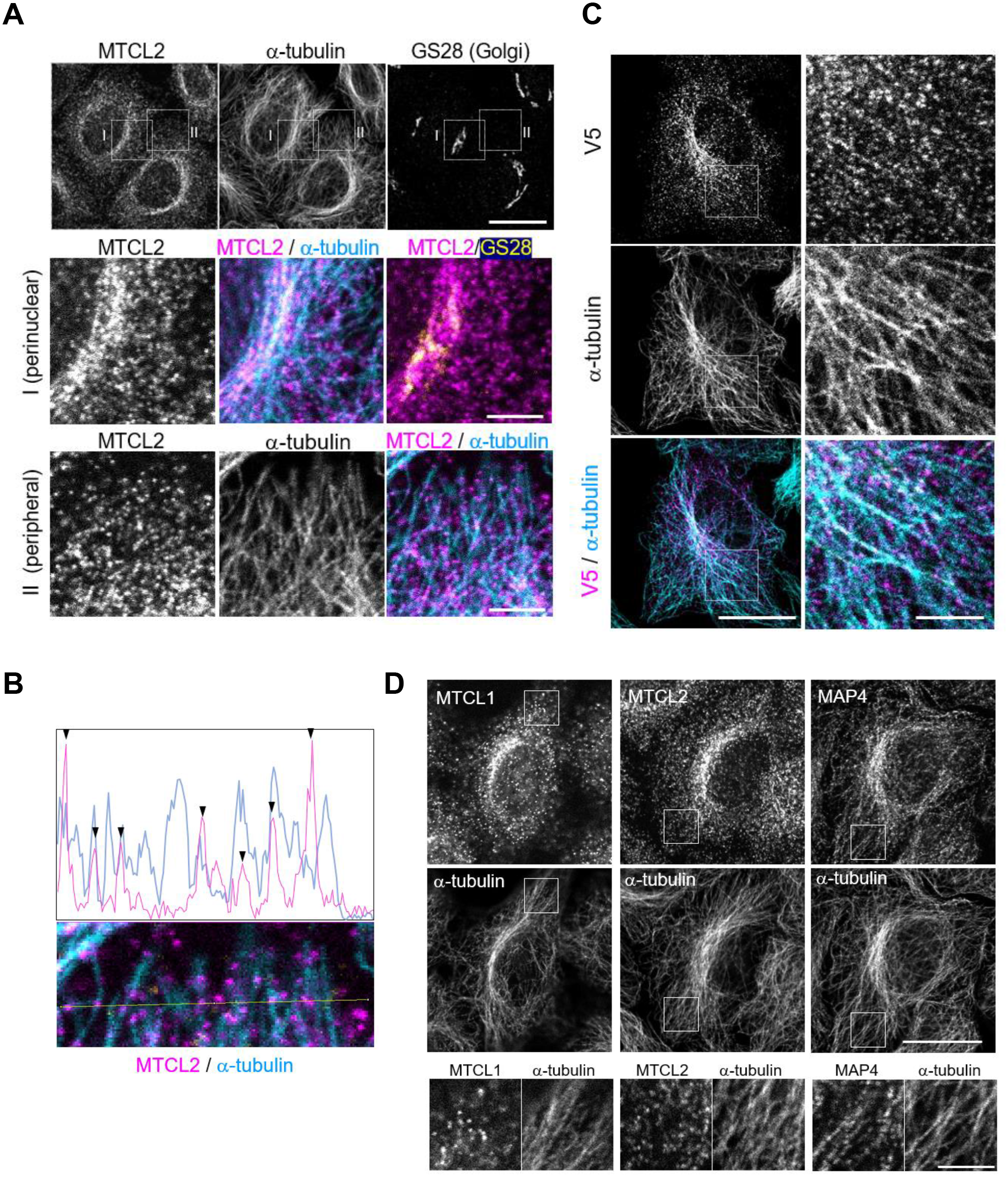
MTCL2 preferentially colocalizes with the perinuclear microtubules accumulated around the Golgi complex. A. HeLa-K cells were stained with anti-SOGA1 (MTCL2) together with anti-α-tubulin and anti-GS28 antibodies. Scale bar: 20 μm. Boxed regions in the top panels I and II are enlarged in middle or bottom panels, respectively. Scale bars: 5 μm. B. Colocalization of MTCL2 with MTs in the peripheral region shown in (A) was confirmed through line scan analysis. C. Subcellular localization of exogenously expressed V5-tagged MTCL2 in HeLa-K cells was analyzed using anti-V5 and anti-α-tubulin antibodies. Scale bar: 20 μm. The boxed region is enlarged in the right panels. Scale bar: 5 μm. D. HeLa-K cells were stained with anti-MTCL1, SOGA1 (MTCL2), or MAP4 antibodies together with anti-α-tubulin antibody, as indicated. Scale bar: 20 μm. Boxed regions are enlarged in bottom panels. Scale bar: 5 μm.

Western blotting analyses of various mouse tissue extracts also revealed a similar ∼200 kDa band as a major band in lanes corresponding to the lung, testis, ovary, cerebrum, and cerebellum (Fig. 1D). Alternatively, weak signals around 80 kDa were detected for some tissues, such as the pancreas, liver, and muscles (arrowheads). Therefore, we cannot exclude the possibility that MTCL2 is subjected to the reported cleavage and functions as a SOGA in these tissues. However, our results are consistent with the notion that MTCL2 is a functional paralog of MTCL1 and ubiquitously serves as an MT-regulating protein in the cytosol.

### MTCL2 predominantly localizes to the perinuclear MTs accumulating around the Golgi complex

Next, we examined the subcellular localization of MTCL2 in HeLa-K cells. Reaction with the anti-SOGA1 antibody yielded granular signals in the cytoplasm (Fig. 2A), which were particularly condensed near the perinuclear region where the Golgi ribbons are located and MTs accumulate (inset I in top panels Fig. 2A). They completely disappeared in MTCL2*-*knockdown cells (Fig. 5), and the same staining patterns were obtained independent of the fixation conditions (Fig. EV2). These results indicate that the immunostaining signals observed herein revealed the genuine localization of endogenous MTCL2. Further analysis indicated that most MTCL2 signals in this perinuclear region were detected on MTs, and some overlapped with the Golgi marker signals (middle panels in Fig. 2A). Colocalization of MTCL2 with MTs was also observed in the peripheral regions (inset II in top panels), where the densities of MTCL2 signals were rather low (bottom panels in Fig. 2A and Fig. 2B).

To further confirm the above results, we examined the subcellular localization of exogenously expressed MTCL2 using an anti-tag antibody. When highly expressed in HeLa-K cells, exogenous MTCL2 induced the formation of thick MT bundles and frequently disrupted the normal crescent-like Golgi ribbon structures into dispersed ones (arrows in Appendix Fig. S1). However, when the expression levels were similar to the endogenous levels, exogenous MTCL2 mimicked endogenous MTCL2 in terms of subcellular localization, by accumulating on one side of the perinuclear region where the Golgi ribbons localize and MTs accumulate (yellow arrowheads in Appendix Fig. S1). Colocalization of exogenously expressed MTCL2 with MTs in the peripheral region was also confirmed in the low-expression conditions (Fig. 2C).

Since the above results were similar to those reported for MTCL1 (Sato *et al*, 2014), we directly compared them (Fig. 2D). As expected, MTCL1 and 2 exhibited a significantly similar distribution pattern in the cytoplasm, with intermittent localization on the MT lattices and preferential condensation on the perinuclear MTs accumulated around the Golgi. These features were highlighted when their distribution patterns were compared with those of another MT lattice-binding protein, MAP4 (Chapin & Bulinski, 1991). Unlike MTCL1 and 2, MAP4 was evenly distributed along MTs, without a strong preference for perinuclear MTs. In the peripheral regions, we could predict directions of each MT filament running in this area from MAP4 signals exhibiting linear arrangements. However, immunofluorescence signals of MTCL1 and 2 were too sparse to enable us to do this prediction (bottom panels in Fig. 2D). These results support the notion that MTCL1 and 2 form a unique family of MT-regulating proteins.

### MTCL2 interacts with MTs via the C-terminal conserved region

To determine the molecular basis of the subcellular localization of MTCL2, we subdivided the molecule into three fragments (N, M, and C in Fig. 3A) and examined their localization in HeLa-K cells (Fig. 3B). As expected, the C fragment containing the region corresponding to MTCL1 C-MTBD (hereafter referred to as the KR-rich region; Fig. 1A, Fig. EV1C) exhibited clear localization on the MT lattice (top and the right panels in Fig. 3B). Direct binding of the C-terminal region with MTs was confirmed using a shorter fragment of MTCL2 (CT1) that still contained the KR-rich region (Fig. 3A): CT1 fused with maltose-binding protein (MBP), but not MBP alone, co-sedimented with MTs *in vitro* when purified from *Escherichia coli* and mixed with taxol-stabilized MTs (Fig. 3C). The KR-rich region alone also exhibited localization on MTs (Fig. 3D, Fig. EV3B), whereas deletion of the KR-rich region impaired localization of full-length MTCL2 on MTs (MTCL2 ΔKR in Fig. EV3A). Together with the results that the N and M fragments did not colocalize with MTs (Fig. 3B), these results indicate that MTCL2 has a single MT-binding region at the C-terminus, as predicted from the sequence comparison between MTCL1 and 2 (Fig. EV1).

**Figure 3.**
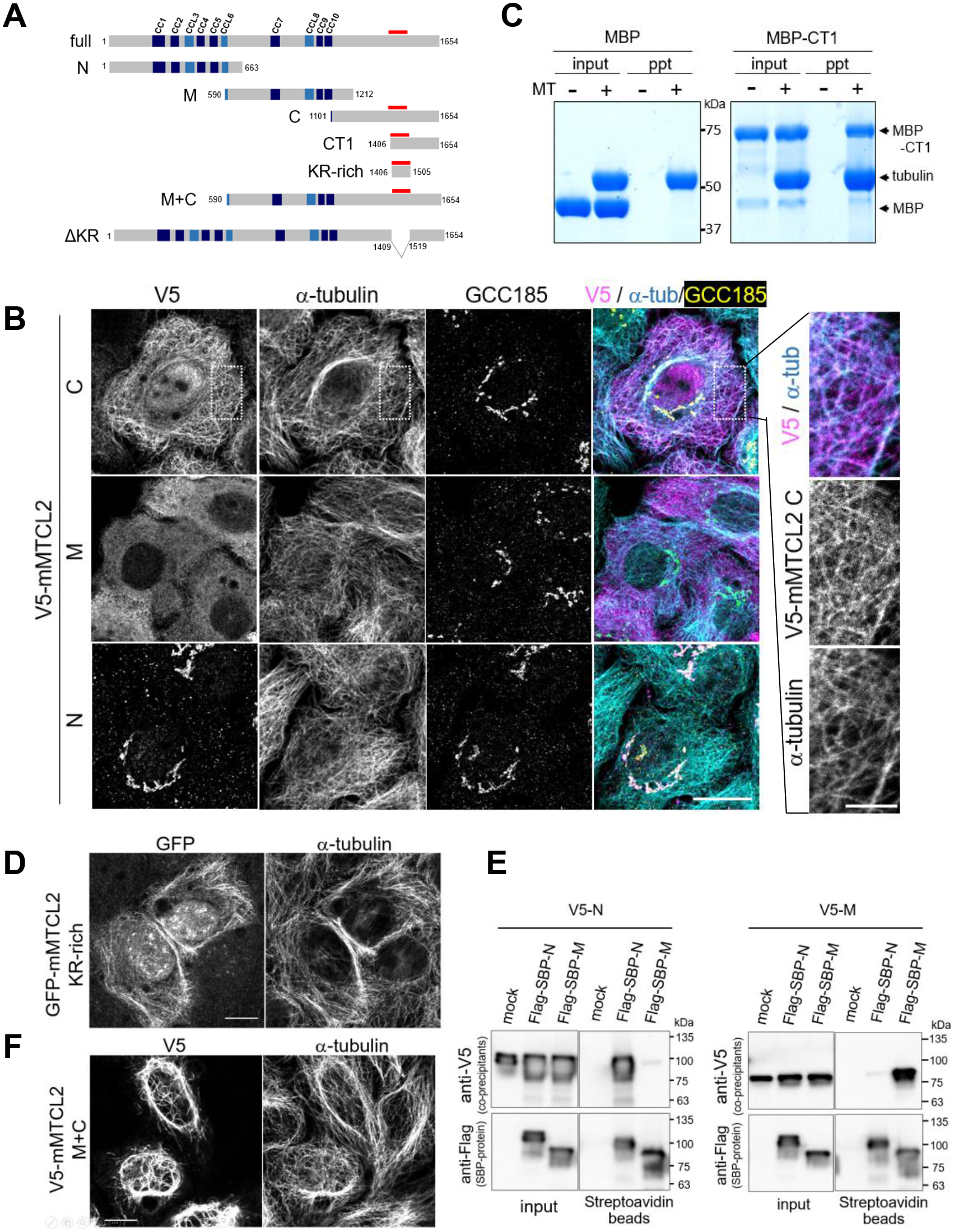
MTCL2 directly associates with MTs via the C-terminal KR-rich region. A. MTCL2 deletion mutants related to this figure. Red bars indicate the position of the KR-rich region. B. Subcellular localization of V5-C (top), -M (middle), and -N (bottom) in HeLa-K cells. The antibodies used are indicated at the top. Scale bar: 20 μm. The boxed region of a V5-C-expressing cell is enlarged in the right panels. Scale bar: 5 μm. Note that the C fragment is colocalized with MTs, whereas the N fragment is localized to the Golgi distinctly. C. MBP-fused CT1 purified from *Escherichia coli* was examined for MT pull-down experiments. MBP-CT1 and not MBP was precipitated only when taxol-stabilized MTs were included. ppt represents the MT precipitate obtained after centrifugation (200,000 × *g*) for 20 min at 25°C. D. Subcellular localization of GFP-KR-rich in HeLa-K cells. Scale bar: 20 μm. E. V5-N (left panels) or V5-M (right panels) were expressed in HEK293T cells together with the indicated proteins and subjected to pull-down assays using streptavidin-conjugated beads. In the mock sample, an empty backbone vector for Flag-SBP constructs was co-transfected. F. Subcellular localization of V5-M+C in HeLa-K cells. Scale bar: 20 μm.

We have previously shown that the C-MTBD of MTCL1 has MT-stabilizing activity (Sato *et al*, 2014; Abdul Kader *et al*, 2017). This activity can be monitored by its ability to strongly increase acetylated tubulin signals as well as to secondarily induce MT bundles when expressed in HeLa-K cells (Fig. EV3B). We noticed that the KR-rich region of MTCL2 did not show these activities strongly (Fig. EV3B). These results suggest that the sequence divergence from MTCL1 (Fig. EV1C) weakened the MT-stabilizing activity of the MT-binding region of MTCL2 and made it similar to MTCL1 N-MTBD, which induced MT bundles only when it was connected to the central coiled-coil region (Abdul Kader *et al*, 2017). To assess this possibility, we first examined homo-interaction of the coiled-coil region of MTCL2 by using N and M fragments tagged with streptavidin-binding peptide (SBP) or V5 peptide. When the fragments with a different tag were expressed in HEK293T cells in various combinations, homo-but not hetero-interactions were detected for the N and M fragments in pull-down experiments using streptavidin-conjugated resin (Fig. 3E). This finding indicated that the central coiled-coil region of MTCL2 mediated parallel oligomerization of MTCL2, similar to MTCL1. Figure 3F demonstrates that the C fragment expressed in HeLa-K cells acquired strong MT-bundling activity when fused with the M fragment. These results support the notion that MTCL2 mainly functions as an MT crosslinking protein by directly interacting with MTs via the C-terminus and forming parallel oligomers via the central coiled-coil region.

### MTCL2 associates with the Golgi apparatus via the N-terminal coiled-coil region

As for MTCL1, we failed to identify the region responsible for its Golgi association activity (unpublished results). However, unexpectedly, a strong association between the N fragment of MTCL2 and the Golgi membrane was observed (bottom panels in Fig. 3B). This finding contrasted sharply with the results that the M fragment distributed diffusely without showing any discrete localizations by itself (middle panels in Fig. 3B), suggesting that MTCL2 is associated with MTs and the Golgi membrane separately through the C- and N-terminal regions, respectively. Considering that the C fragment did not exhibit preferential localization to the perinuclear region (top panels in Fig. 3B), this dual binding activity of MTCL2 may enable the exhibition of preferential association with the perinuclear MTs around the Golgi apparatus.

To identify mutations that disrupt the Golgi association of the N fragment, we first performed deletion mapping of a region responsible for this Golgi-association activity and found that the most N-terminal region highly diverged from MTCL1 was dispensable (NΔN in Fig. 4A, Fig. EV1). However, subsequent analysis did not allow us to confine the responsible region narrower than 431 amino acids covering the six N-terminal CC motifs (CC1–CCL6) and an additional ∼40 amino acid sequence downstream of CCL6 named Golgi-localizing essential domain (GLED] (CC1-GLED; Fig. 4A, Appendix Fig. S2). We then examined the effects of point mutations in the coiled-coil motifs of the N fragment. At first, four leucine residues appearing in every seven amino acids in the first half of CC1 were mutated to proline (4LP) to disrupt the α-helix itself, or alanine (4LA) to preserve the α-helical structure but suppress its hydrophobic interactions (Fig. 4C). Importantly, not only 4LP mutation but also 4LA mutation was found to be sufficient to disrupt the Golgi localization of the N fragment (Fig. 4D). These results indicate that the coiled-coil interaction through the first half of CC1 was crucial for the Golgi association of the N fragment. We confirmed that 4LA mutations did not disrupt the co-assembling activity of the N fragment (Appendix Fig. S3), likely owing to the homo-interaction of the remaining coiled-coil motifs. This finding indicates that a partial disturbance of the oligomerization state of the N fragment was sufficient to disrupt the Golgi association.

**Figure 4.**
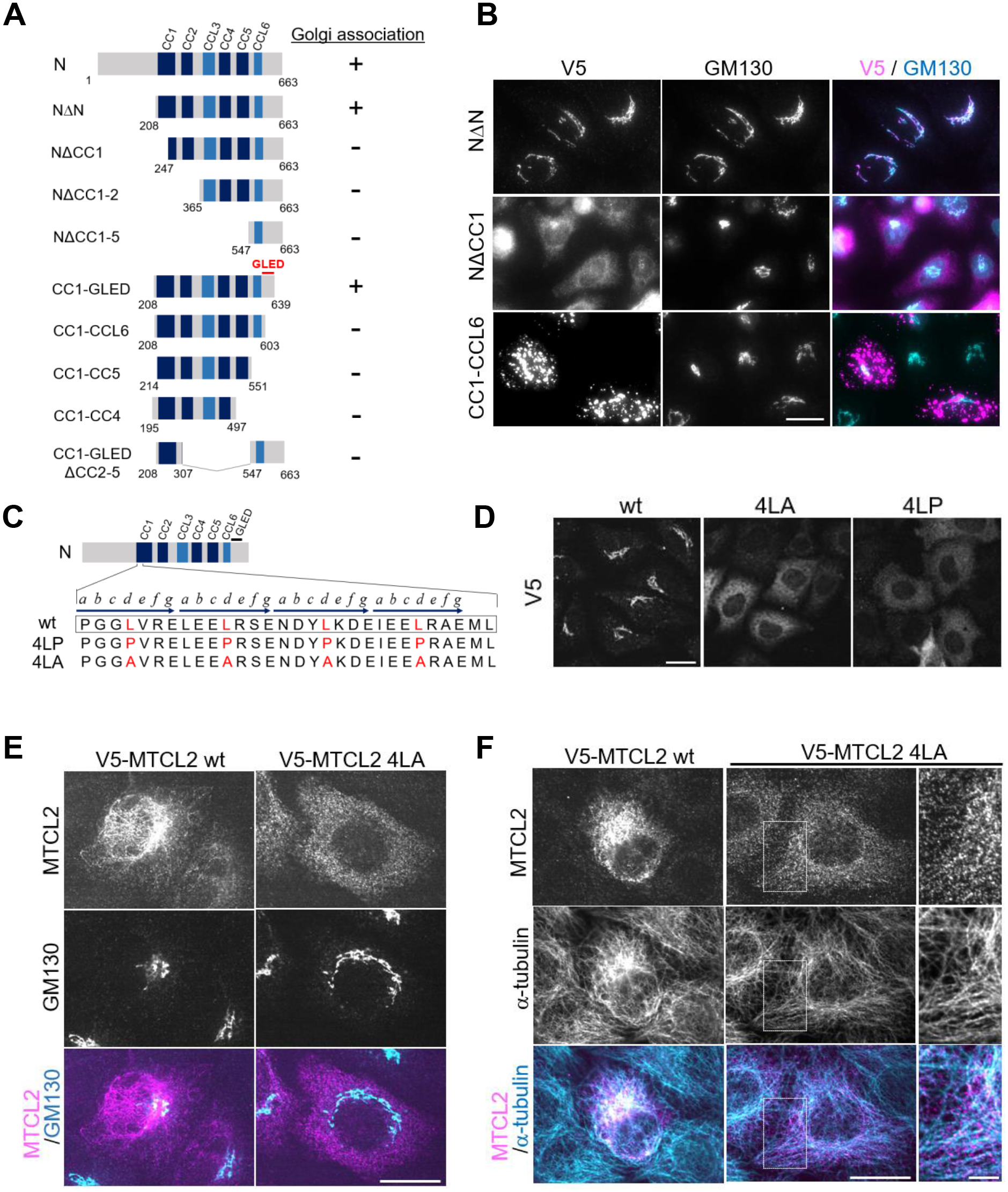
MTCL2 associates with the Golgi membrane via the N-terminal coiled-coil region. A. MTCL2 deletion mutants related to this figure and the summary of their Golgi association activity. B. Subcellular localization of the indicated mutants in HeLa-K cells. Scale bar: 20 μm. C. Amino acid sequence of the first half of MTCL2 CC1 and positions of four leucine residues mutated in this study (red). The characteristic seven-residue repeats are indicated by horizontal arrows, and the positions of each amino acid in a repeat are indicated by italic alphabets (*a* to *g*). D. Subcellular localization of the indicated N fragment mutants in HeLa-K cells was examined using anti-V5 antibody. Scale bar: 20 μm. E and F. Subcellular localization of V5-mMTCL2 wt or 4LA mutant in HeLa-K cells analyzed using anti-MTCL2 (SOGA1) antibody together with anti-GM130 (E) or anti- α-tubulin (F). Scale bar: 20 μm. Boxed regions in (F) are enlarged in the right panels. Scale bar: 5 μm. Note that the expression of each exogenous protein was induced in MTCL2-knockdown cells and suppressed to the endogenous level.

Next, we examined whether these mutations affected the subcellular localization of full-length MTCL2. In these experiments, the expression of exogenous MTCL2 was induced at the endogenous levels in MTCL2-knockdown cells to exclude the effect of endogenous MTCL2 (Materials and Methods). In contrast to wild-type MTCL2, which showed preferential localization to the perinuclear MTs, similar to endogenous MTCL2, the 4LA mutant was diffusely distributed in the cytoplasm without any condensation around the Golgi (Fig. 4E). Importantly, careful examination revealed its colocalization with MTs (Fig. 4F, Fig. EV3A), suggesting that MTCL2 can interact with MTs independent of its Golgi association. These findings indicate that the characteristic perinuclear accumulation of endogenous MTCL2 was the result of its Golgi association through the N-terminal coiled-coil region.

### MTCL2 promotes the accumulation of MTs around the Golgi complex by crosslinking MTs on the Golgi membrane

We analyzed the effects of MTCL2 knockdown in HeLa-K cells to explore the physiological function of MTCL2. For this purpose, we first established heterogeneous stable cells expressing mMTCL2 in a doxycycline-dependent manner (Materials and Methods). When cells were transfected with control siRNA in the absence of doxycycline (without exogenous MTCL2 expression), normal accumulation of MTs around the perinuclear region at which endogenous MTCL2 was concentrated was observed (Fig. 5A). Alternatively, when cells were subjected to MTCL*2* knockdown in the absence of doxycycline (without exogenous MTCL2 expression), MT accumulation around the perinuclear region was severely reduced (Fig. 5A). The specificity of these knockdown effects was confirmed by a rescue experiment in which doxycycline was added to induce the expression of RNAi-resistant MTCL2 (mMTCL2) at endogenous levels. Under these conditions, many cells restored MT accumulation in the perinuclear region, where exogenous MTCL2 was concentrated. We quantitatively estimated the asymmetric distribution of MTs by calculating the skewness of the intensity distribution of tubulin signals within each cell (Fig. 5B, Appendix Fig. S4). In the control cells, the pixel intensity of tubulin signals was distributed with a skewness of 1.02 (median), whereas in MTCL2-knockdown cells, this value decreased to 0.73, indicating a more symmetric distribution of MTs. The expression of RNAi-resistant mMTCL2 restored this value to 1.17, statistically supporting its rescue activity.

**Figure 5.**
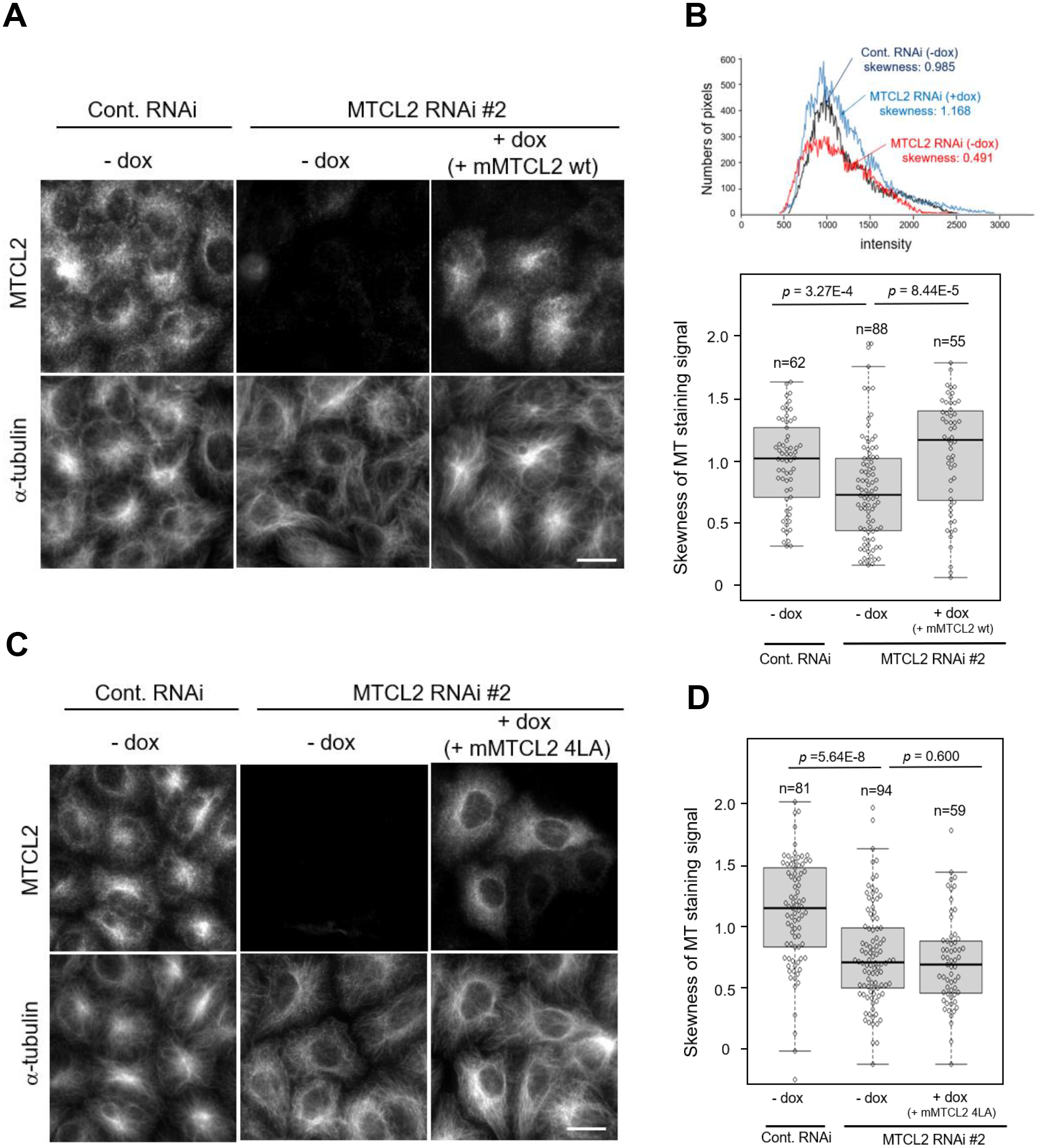
MTCL2 promotes the perinuclear accumulation of MTs in a Golgi-association-dependent manner. A. HeLa-K cells stably harboring pOSTet15.1 expression vector for mouse MTCL2 were transfected with siRNAs for control or MTCL2 knockdown (#2) in the presence or absence of 100 nM doxycycline (dox) and doubly stained with anti-SOGA1 (MTCL2) and anti-α-tubulin antibody, as indicated on the left. Scale bar: 20 μm. B. Extent of MT accumulation was quantitatively estimated by calculating the skewness of the pixel intensity distribution for tubulin signals in each cell. The top panel shows typical data on the tubulin signal distributions and their skewness values, indicating that the asymmetries of tubulin signal distribution are compromised in MTCL2-knockdown cells. The bottom is a box plot of the skewness distribution in each condition. The lines within each box represent medians. Data represent the results of the indicated number (n) of cells from a typical experiment (biological replicates; Materials and Methods). The *p* values were estimated using the Wilcoxon test. Statistical data of technical replicates (three independent experiments) are demonstrated in Appendix Fig. S4. C. HeLa-K cells stably harboring pOSTet15.1 expression vector for mouse MTCL2 4LA were subjected to the same experimental procedure as in (A). Scale bar: 20 μm. D. MT accumulation data in (C) was quantitatively analyzed as in (B).

Interestingly, MTCL2 knockdown also affected the assembly structure of the Golgi stacks (Fig. EV4A). In contrast to control cells, which showed a compact crescent-like morphology of the Golgi ribbon on one side of the nucleus, MTCL2-knockdown cells exhibited abnormally expanded Golgi ribbons along the nucleus (Fig. EV4A). The median expansion angle (θ) of the Golgi apparatus (Fig. EV4A) was 65.4° for the control cells, whereas it significantly increased to 82.5° in MTCL2-knockdown cells (Fig. EV4B, Appendix Fig. S4). The expression of RNAi-resistant MTCL2 reduced the angle with a median value of 61.0°, indicating that MTCL2 was essential for compact accumulation of the Golgi ribbon. Similar effects of MTCL2 knockdown were observed in RPE1 cells (Appendix Fig. S5). These results demonstrate that MTCL2 plays an important role in promoting the perinuclear accumulation of MTs and increasing the compactness of Golgi ribbons.

Considering the MTCL2 activities shown in Figs. 2 and 3, these results are highly consistent with the hypothesis that MTCL2 crosslinks MTs on the Golgi membrane, thereby accumulating MTs around the Golgi ribbon. The effects on the compactness of the Golgi ribbon can also be explained as a secondary effect of MT accumulation, which must attract individual Golgi stacks to each other (Fig. EV4C). To test this hypothesis, we performed the same experiments using stable cells expressing the 4LA mutant in a doxycycline-dependent manner (Fig. 5C and D, Fig. EV4D and E, Appendix Fig. S4). Knockdown effects on MT organization and Golgi ribbon compactness were similarly observed in these stable cells (-dox). However, the expression of the 4LA mutant (+dox) did not restore both phenotypes. These findings indicate the importance of Golgi association in MTCL2 functioning. Through similar experiments, we further confirmed that MTCL2 lacking the MT-binding region (MTCL2 ΔKR) also showed loss of rescue activities against both phenotypes (Fig. EV5, Appendix Fig. S4).

Altogether, we conclude that MTCL2 promotes MT accumulation around the Golgi ribbon by exerting its MT crosslinking activity on the Golgi membrane.

### MTCL2 depletion resulted in defects in cell migration

The Golgi ribbon structure and its associated MTs are essential for maintaining directed cell migration owing to their essential roles in the polarized transport of vesicles (Bergmann *et al*, 1983; Yadav *et al*, 2009b; Miller *et al*, 2009; Sato *et al*, 2014; Hurtado *et al*, 2011). Therefore, we next examined whether MTCL2 depletion affected directed cell migration during the wound healing process *in vitro*.

First, HeLa-K cells transfected with control or MTCL2 siRNA were grown to a confluent monolayer and scratched with a micropipette tip to initiate directional migration into the wound. In control cells at the wound edge, reorientation of the Golgi and elongation of a densely aligned MT toward the wound were observed (Fig. 6A). In MTCL2-knockdown cells, reorientation of the Golgi was reduced but not severely affected. Nevertheless, cells lacking MTCL2 exhibited randomly oriented MTs and failed to align them toward the wound (Fig. 6A).

**Figure 6.**
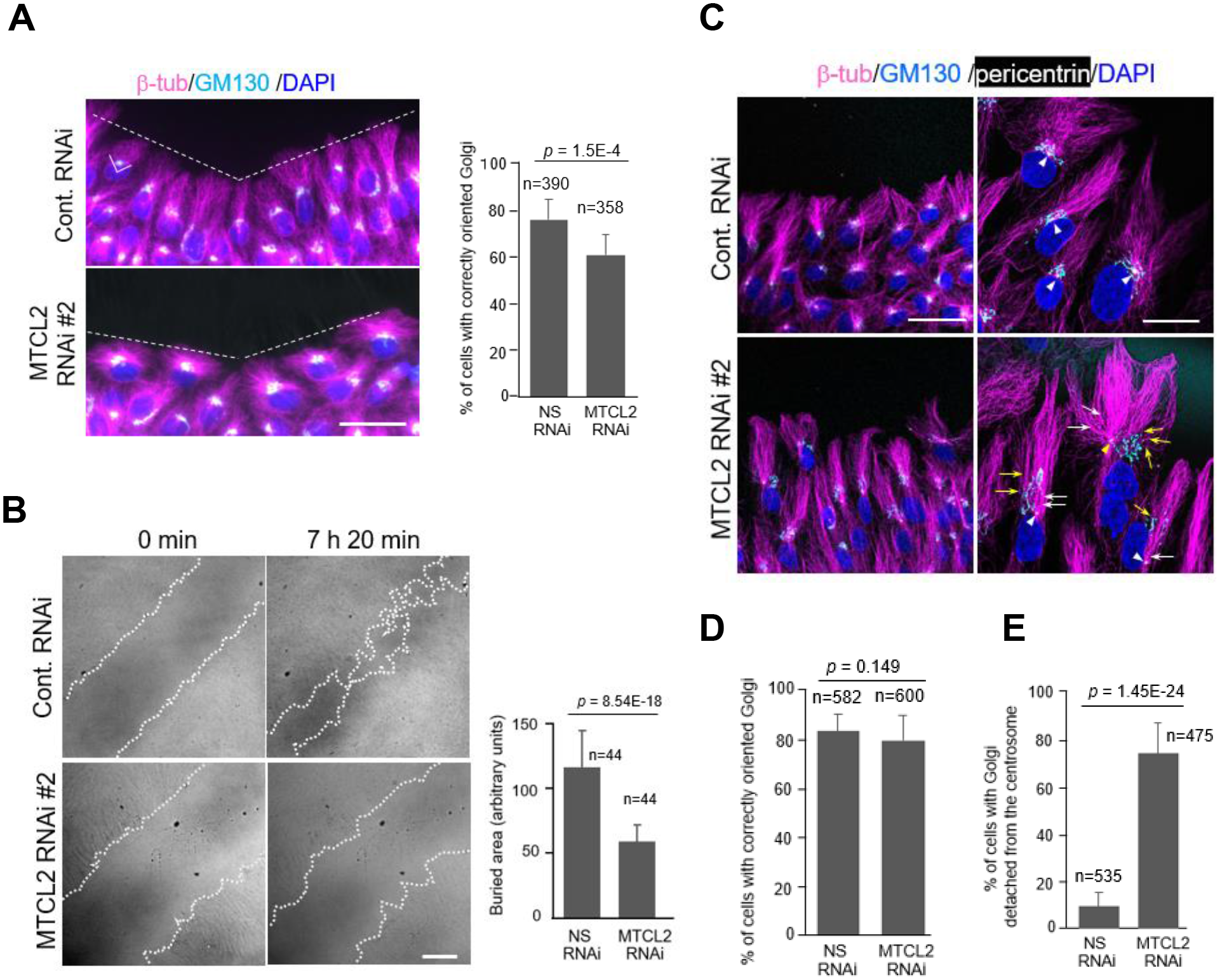
MTCL2 depletion results in defective cell migration. A. Confluent monolayers of HeLa-K cells subjected to control or MTCL2 RNAi were fixed and stained with the indicated antibodies 6 h after wounding. Cells facing the wound edges (white dotted lines) are shown. Scale bar: 50 μm. Note that MTCL2-depleted cells did not polarize MT arrays toward the wound. The right panel indicates the percentage of wound-edge cells with correctly oriented Golgi, defined as those falling in the indicated quadrant (white line) concerning the wound edge. Data represent the means ± S.D. for the indicated number (n) of cells from two independent experiments. The *p* value was estimated using the Student’s t-test assuming the two-tailed distribution and two-sample unequal variance. B. Differential interference contrast images of wound healing RPE1 cells at 0 min and 7 h 20 min after wounding. White dotted line delineates the wound edges. Scale bar: 200 μm. Right panel indicates quantified data on the areas newly buried by cells after wounding. Data represent the mean ± S.D. of 44 fields taken from two independent experiments. The *p* value was estimated using Student’s t-test assuming the two-tailed distribution and two-sample unequal variance. C. RPE1 cells subjected to wound healing analysis in (B) were fixed and stained with the indicated antibodies. Cells facing the wound edges are shown. Right panels show the enlarged view. Arrowheads indicate the positions of the centrosomes. Note that MTCL2-depleted cells exhibit separation of the centrosome and Golgi. The centrosomes frequently show significant detachment from the perinuclear region (see yellow arrowhead). White and yellow arrows indicate MTs emanating from the centrosome and the Golgi, respectively. Scale bars: 50 μm and 20 μm (enlarged right panels). D. Golgi orientation was quantified for wound healing RPE1 cells, as indicated in (A). Data represent the means ± S.D. for the indicated number (n) of cells from two independent experiments. The *p* value was estimated using the Student’s t-test assuming the two-tailed distribution and two-sample unequal variance. E. Percentage of wound-edge cells with Golgi detached from the centrosome. Data represent the means ± S.D. for the indicated number (n) of cells from two independent experiments. The *p* value was estimated using the Student’s t-test assuming the two-tailed distribution and two-sample unequal variance.

Despite the significant difference in MT organization in cells at the wound edge, we could not estimate the effects of MTCL2 knockdown on directional migration as the HeLa-K cells migrated very slowly. Thus, we used RPE1 cells to estimate wound healing velocity and found that cells lacking MTCL2 migrated significantly slower than control cells (Fig. 6B, Movies EV1 and 2). Comparison of the normalized areas newly covered by migrated cells revealed that the directed migration velocity of MTCL2-knockdown cells was approximately 50% of that of control cells (Fig. 6B, right panel). Time-lapse analysis of differential interference contrast images indicated that cells lacking MTCL2 exhibited abnormally elongated shapes and were less efficient in extending lamellipodia (Movie EV2). Reorientation of the Golgi position toward the wound was observed in MTCL2-knockdown cells to a similar extent as in control cells (Fig. 6C, D). In addition, in contrast to HeLa-K cells, MTCL2-knockdown cells showed polarized elongation of MTs toward the wound (Fig. 6C). However, the proximal ends of these MTs seemed unfocused. Close inspection revealed that in MTCL2-knockdown cells at the wound edge, the Golgi ribbon was frequently separated from the centrosome and sometimes detached from the nucleus (Fig. 6C and E). As a result, the centrosomal MTs and Golgi-associated MTs elongated from different positions toward the wound and were discerned in many MTCL2-knockdown cells (arrows in Fig. 6C, right panel). By contrast, in control cells, the centrosome and Golgi ribbon were tightly linked near the nucleus, and the proximal ends of the centrosomal and Golgi-associated MTs were indistinguishable. These data suggest an intriguing possibility that MTCL2 may play an essential role in integrating centrosomal and Golgi-associated MTs by crosslinking them on the Golgi membrane.

### CLASPs are required for the perinuclear localization of MTCL2

The Golgi association of MTCL1 was shown to be mediated by CLASPs and AKAP450 (Sato *et al*, 2014). Therefore, to identify proteins that mediate the Golgi association of MTCL2, we first examined the effect of knockdown of CLASP or AKAP450 on the subcellular localization of MTCL2. Simultaneous depletion of CLASP1 and 2 profoundly impaired the accumulation of MTCL2 in the perinuclear region and induced an even cytosolic distribution (Fig. 7A, Appendix Fig. S6). AKAP450 depletion also affected the distribution of MTCL2; however, it did not induce dissociation of MTCL2 from the perinuclear region where the Golgi apparatus localizes (Fig. 7A, Appendix Fig. S6). These results are consistent with the report that SOGA1 interacts with CLASP2 (Kruse *et al*, 2017) and suggest the possibility that CLASPs play major roles in mediating the Golgi association of MTCL2. Consistently, GFP-CLASP2α specifically interacted with N but not the M or C fragment of MTCL2 when co-expressed in HEK293T cells and subjected to pull-down experiments (Fig. 7B). The interaction was also observed for the minimum fragment of MTCL2 (CC1-GLED) required for Golgi association (Fig. 7B). However, we unexpectedly observed substantial interactions between GFP-CLASP2α and CC1-GLED with 4LA mutations within CC1. In addition, depletion of AKAP450 and CLASPs did not affect the Golgi localization of the N fragment exogenously introduced in HeLa-K cells (Fig. 7C). These results raised the possibility that unknown factors other than CLASPs were involved in the CC1-dependent interaction of MTCL2 with the Golgi membrane. To identify these putative factors, we screened Golgi marker proteins exhibiting the most precise colocalization with the N fragment of MTCL2 (Appendix Fig. S7A). Close inspection using super-resolution microscopy revealed that the N fragment showed distinct localization from cis- and trans-Golgi markers; however, it exhibited the most significant colocalization with a cis/medial marker, giantin/GOLGB1 (Appendix Fig. S7A) (Linstedt *et al*, 1995). This finding led us to find that the Golgi localization of the N fragment almost disappeared in cells lacking giantin (Fig. 7C). Since expression of the N fragment was not reduced in giantin-knockdown cells (Appendix Fig. S7B and C), these results indicated that giantin is required for the Golgi association of the MTCL2 N-terminus. Giantin knockdown partially impaired the perinuclear accumulation of endogenous MTCL2 (Fig. 7A, Appendix Fig. S6). Collectively, the findings indicate the possibility that giantin is primarily responsible for the recruitment of MTCL2 to the Golgi membrane in a CC1-dependent manner before CLASP involvement. As endogenous MTCL2 only shows restricted colocalization with CLASPs or giantin (Appendix Fig. S8), the interactions between MTCL2 and these proteins might be gradually weakened to realize its steady-state localization predominantly associated with the perinuclear MTs.

**Figure 7.**
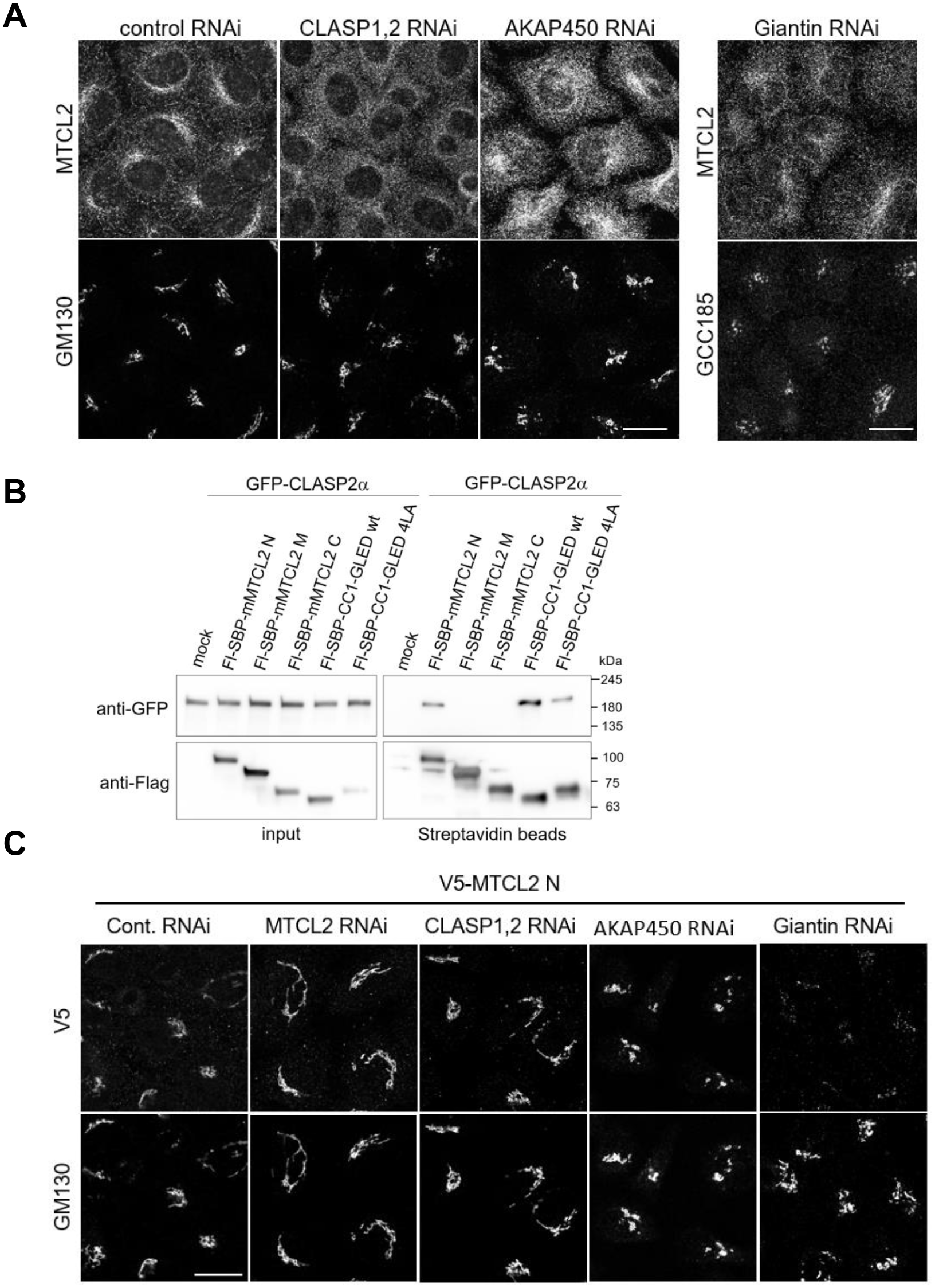
CLASPs and giantin are involved in the Golgi association of MTCL2. A. HeLa-K cells subjected to the indicated knockdown were stained with anti-SOGA1 (MTCL2) and anti-GM130 antibodies. Scale bar: 20 μm. B. GFP-CLASP2α was expressed in HEK293T cells together with the indicated Flag-SBP-tagged proteins and subjected to pull-down assays using streptavidin-conjugated beads. C. V5-N was expressed in HeLa-K cells subjected to the indicated knockdown and examined for the Golgi association. Scale bar: 20 μm.

## Discussion

The results of this study demonstrate that the MTCL1 paralog MTCL2 is a novel MT-regulating protein that preferentially associates with perinuclear MTs around the Golgi. Its dual binding activity to MTs and the Golgi, as well as its oligomerizing activity via the coiled-coil region, collectively enabled it to crosslink and accumulate MTs on the Golgi membrane. Our data suggest that this unique activity of MTCL2 plays an important role in directed migration by integrating the centrosomal and Golgi-nucleated MTs on the Golgi membrane.

MTCL2 depletion severely disrupted the accumulation of MTs around the Golgi and induced random arrays of MTs (Fig. 5A). Low-dose re-expression of MTCL2 restored the original organization of MTs in a Golgi-binding activity-dependent manner. These data indicate that MTCL2 plays an indispensable role in the asymmetric organization of global MTs by utilizing the Golgi complex as a foothold for its MT-crosslinking function (Meiring *et al*, 2020). These findings also highlight the active role of the Golgi complex in MT organization in interphase cells. Regarding the molecular mechanisms underlying the Golgi association of MTCL2, we provide data indicating the possible involvement of CLASPs and giantin. CLASPs have been shown to play essential roles in development of Golgi-associated MTs through its +Tips activity (Miller *et al*, 2009; Efimov *et al*, 2007). Our results indicate that MT regulation by CLASPs is also mediated by its novel activity to facilitate the perinuclear condensation of MTCL2 (Appendix Fig. S6B). The present data also reveal the presence of complicated mechanisms in the Golgi association of MTCL2, because CLASP2 interacted with the Golgi association region of MTCL2 independent of the 4LA mutation, and was not necessarily required for its recruitment to the Golgi membrane. These complexities are consistent with the requirement of a long amino acid sequence containing multiple coiled-coil motifs and an additional unstructured region (GLED) (Appendix Fig. S2B) for the Golgi association, suggesting the presence of multiple and sequential interactions between the MTCL2 N-terminus and several Golgi-resident proteins. Here, we demonstrated that one of the candidate molecules of these Golgi-resident proteins is giantin, a large coiled-coil protein, the involvement of which has been demonstrated in ER-Golgi traffic as a tethering factor (Sönnichsen *et al*, 1998; Alvarez *et al*, 2001). The complexity of the Golgi-binding mechanisms is also indicated by comparison with MTCL1, which also exhibits a subcellular localization strikingly similar to that of MTCL2 (Fig. 2D) (Sato *et al*, 2014). As for this MTCL protein, we have failed to identify the Golgi association region, despite the significant amino acid similarity of its N-terminal coiled-coil motifs with MTCL2 (Fig. EV1). This difference between these paralogs is not merely due to the fact that the GLED sequence of MTCL2 is not conserved in MTCL1 (Fig. EV1B). The seamless exchange of the highly conserved CC1 sequence between MTCL1 and 2 was sufficient to disrupt the Golgi localization of the N fragment of MTCL2 (Appendix Fig. S2C). This finding indicates that the Golgi-binding mechanisms of MTCL proteins are not based on simple coiled-coil interactions but consist of sophisticated protein–protein interactions that are highly differential between these paralogues. Actually, we found here that the Golgi association of MTCL2 strongly depends on CLASPs but not AKAP450 (Fig. 7A), whereas the Golgi association of MTCL1 depends on both proteins almost evenly (Sato *et al*, 2014). In this context, it is also noteworthy that full-length MTCL2 lacking MT-binding activity (MTCL2 DKR) was distributed diffusely without Golgi localization (Fig. 3A, Fig. EV3A). This finding contrasted sharply with the clear association of the N fragment with the Golgi membrane (Fig. 3B). These results indicate a possibility that MT binding through the C-terminal region is a prerequisite for Golgi association via the N-terminal coiled-coil region and imply intramolecular regulation of the Golgi binding of MTCL2. The fact that endogenous MTCL2 does not exhibit complete colocalization with the Golgi complex further suggests the presence of additional mechanisms that regulate the balance between the dual binding activities of MTCL2 to MTs and the Golgi membrane. Our results indicate that MTCL2 is expressed in several cultured cells simultaneously with MTCL1 (Fig. 1C) (Sato *et al*, 2014). Tissue distribution patterns of MTCL2 were also found to be similar to that of MTCL1 (Fig. 1D) (Satake *et al*, 2017). These results raise questions regarding how cells utilize these MTCL proteins. A clue can be drawn from the previously reported result that, in contrast to MTCL2, MTCL1 knockdown does not change the global organization of MTs significantly but only reduces a specific subpopulation of MTs specifically stained with an anti-acetylated tubulin antibody (Sato *et al*, 2014). This MT subpopulation corresponds to stable MTs that are nucleated from the Golgi membrane with the aid of CLASPs and AKAP450 (Chabin-Brion *et al*, 2001; Rivero *et al*, 2009; Efimov *et al*, 2007). We have demonstrated that MTCL1 stabilizes and crosslinks this specific MT subpopulation via C-MTBD and N-MTBD, respectively (Sato *et al*, 2014; Abdul Kader *et al*, 2017). Collectively, these results suggest that the target of MTCL1 action is restricted to the Golgi-associated MTs. Alternatively, we observed here that MTCL2 knockdown dramatically changed the global organization of MTs (Fig. 5A), and that the MT-binding region of MTCL2 lacks strong activity to stabilize MTs (Fig. EV3). These results suggest the possibility that, in contrast to MTCL1, the target of MTCL2 action may not be restricted to the Golgi-nucleated MTs. In extreme cases, MTCL2 may crosslink any kinds of MTs running nearby the Golgi complex. This idea appears to be consistent with the present observation that MTCL2 is required to integrate centrosomal and Golgi-derived MTs on the Golgi membrane. Distinct involvement of AKAP450 in Golgi recruitment might be one of the bases of these functional differences between MTCL1 and 2. Further assessment of the Golgi recruiting mechanisms of each protein will better elucidate this issue.

In this study, we established the presence of a new family of MT-regulating proteins, the MTCL family. Since the members of this family are only found in vertebrates, their functions are expected to be tightly linked with vertebrate-specific cellular structures and functions. We propose to call this gene product MTCL2 instead of SOGA1 because our results demonstrate that it is a functional homolog of MTCL1 in the full-length form and does not function as SOGA1 in a cleaved form ubiquitously. This claim is also based on the fact that we failed to confirm the presence of a putative internal signal sequence as well as Atg16- and Rab5-binding motifs in the MTCL2 sequence, all of which have been discussed previously (Cowherd *et al*, 2010).

## Materials and Methods

### Vector production

The cDNA clone encoding full-length mMTCL2 (GenBank accession number: AK147227) was purchased from Danaform (Kanagawa, Japan). After confirming sequence identity with NM_001164663, a DNA fragment corresponding to the MTCL2 open reading frame was subcloned into an expression vector, pCAGGS-V5. Subsequently, several deletion mutants of MTCL2 were constructed in pCAGGS-V5, pEGFP-c2 (Takara Bio Inc., Japan), or pMal-c5x (New England Biolabs). To establish heterogeneous stable transformants, mMTCL2 and its mutants with or without a 6 × V5-tag were subcloned in pOSTet15.1 (kindly provided by Y. Miwa, University of Tsukuba, Japan), an Epstein–Barr virus-based extrachromosomal vector carrying a replication origin and replication initiation factor sufficient for autonomous replication in human cells (Tanaka *et al*, 1999). Mouse MTCL1 cDNA (GenBank accession number: AK147691) was used as described previously (Sato *et al*, 2013). Expression vector for GFP-CLASP2α was a gift from I, Hayashi (Yokohama City University, Japan).

### Antibodies

To detect MTCL1 and MTCL2, anti-KIAA0802 (sc-84865) from Santa Cruz Biotechnology and anti-SOGA1 (HPA043992) from Sigma-Aldrich were used, respectively. To detect other proteins, the following antibodies were used: anti-α-tubulin (sc-32293), anti-acetylated tubulin (sc-23950), anti-MAP4 (sc-67152), anti-GFP (sc-9996) and anti-CLASP2 (sc-376496) from Santa Cruz Biotechnology; anti-V5 (R960-25) from Thermo Fisher Scientific; anti-GM130 (610822) and anti-GS28 (611184) from BD Transduction Laboratories; anti-GAPDH (5G4) from HyTest Ltd.; anti-giantin (ab37266) and anti-pericentrin (ab4448) from Abcam; anti-GCC185 from Bethyl Laboratories; anti-Flag (F3165) from Sigma-Aldrich; anti-β-tubulin (MAB3408) from Merk Millipore; anti-CLASP1 (MAB9736) from Abnova; anti-AKAP450 from Novus Biologicals; anti-Golgin97 (D8P2K) from Cell Signaling Technology.

### Cell culture and plasmid transfection

HeLa-Kyoto (HeLa-K), HEK293T, and HepG2 cells were cultured in Dulbecco’s modified Eagle’s medium (DMEM, low glucose; Nissui, Japan) containing 10% fetal bovine serum, 100 U/mL penicillin, 100 μg/mL streptomycin, and 1 mM glutamine at 37°C in 5% CO_2_. The hTERT-immortalized human retinal pigment epithelial cells (RPE1 cells) were maintained in a 1:1 mixture of DMEM/Ham’s F12 (FUJIFILM Wako Pure Chemical Corporation, Japan) containing 10% fetal bovine serum, 100 U/mL penicillin, 100 μg/mL streptomycin, 10 μg/mL hygromycin B, and 1 mM glutamine at 37°C in 5% CO_2_. For immunofluorescence analysis, cells were seeded on coverslips in 24-well plates and coated with atelocollagen (0.5 mg/mL IPC-50; KOKEN, Japan). Plasmid transfections were performed using polyethyleneimine (Polysciences, Inc.) for HEK293T cells or Lipofectamine LTX (Life Technologies Corporation) for HeLa-K cells according to the manufacturer’s instructions. To establish heterogenous stable HeLa-K cells expressing mMTCL2 or its mutants in a doxycycline-dependent manner, cells were transfected with pOSTet15.1 expression vector encoding the appropriate MTCL2 cDNA. The following day, cells were reseeded at one-twentieth of the cell density and subjected to selection using a medium containing 800 μg/mL G418 disulfate (Nacalai Tesque, Japan) for more than six days. Surviving cells were used in subsequent experiments without cloning.

### RNAi experiments and wound healing assays

siRNAs for human MTCL2 were designed to target the following sequences: MTCL#2, GAGCGACCGAGAGAGCATTCC; #5, CTGAAGTACCGCTCGCTCT. The target sites for CLASP1, CLASP2, and AKAP450 have been described previously. CLASP1, GGATGATTTACAAGACTGG; CLASP2, GACATACATGGGTCTTAGA (Mimori-Kiyosue et al, 2005); AKAP450, AACTTTGAAGTTAACTATCAA (Rivero *et al*, 2009). A non-silencing RNAi oligonucleotide (Allstars negative control siRNA) was purchased from Qiagen. Cells were seeded on coverslips at densities of 1.2–4 × 10^4^ cells and transfected with siRNAs at final concentrations of 10–17 nM using RNAiMax (Life Technologies Corporation) according to the manufacturer’s instructions. siRNA transfection was repeated the day after medium change, and cells were subjected to immunofluorescence analysis on day 3. For rescue experiments, heterogeneous stable HeLa-K cells expressing mMTCL2 were subjected to a similar protocol, except that 100 ng/mL of doxycycline was always included in the medium after the first siRNA transfection. For wound healing analysis, HeLa-K cells subjected to RNAi were scratched with a micropipette tip on day 4. RPE1 cells were seeded at 5 × 10^4^ cells in one compartment of a 35 mm glass bottom culture dish separated into four compartments (Greiner, 627870) after coating with 10 μg/mL fibronectin (FUJIFILM, Japan, 063-05591). The siRNA transfections were performed as described above. Wounds were created on day 4 by scratching the cell monolayers with a micropipette tip and subjected to live imaging.

### Cell extraction and western blotting

Cell extracts were prepared by adding RIPA buffer (25 mM Tris/HCl [pH 7.5], 150 mM NaCl, 1% NP40, 1% deoxycholic acid, 0.1% SDS) containing a protease inhibitor cocktail (Sigma-Aldrich, P8340) followed by brief sonication and centrifugation (15,000 × *g*, 15 min). For tissue distribution analysis of MTCL2, mouse tissue lysates prepared in a previous study were used (Satake *et al*, 2017). Samples were separated by SDS-PAGE and transferred to polyvinylidene fluoride membranes. Blots were incubated in blocking buffer containing 5% (w/v) dried skim milk in PBST (8.1 mM Na_2_HPO_4_.12H_2_O, 1.47 mM KH_2_PO_4_, 137 mM NaCl, 2.7 mM KCl, and 0.05% Tween 20), followed by overnight incubation with the appropriate antibodies diluted in blocking buffer. Dilutions of anti-SOGA1 and anti-GAPDH antibodies were 1:1000 and 1:5000, respectively. The secondary antibodies were diluted at 1:2,000. Blots were then exposed to horseradish peroxidase (HRP)-conjugated secondary antibodies (GE Healthcare) diluted in blocking buffer for 60 min at RT and washed again. Blots were visualized using Immobilon Western Chemiluminescent HRP Substrate (Millipore) or ECL western blotting detection system (GE Healthcare). Chemiluminescence was quantified using the ImageQuant LAS4000 Luminescent Image Analyzer (GE Healthcare).

### Immunofluorescence staining

In most cases, cells were fixed with cold methanol for 10 min at −20°C, followed by blocking with 10% (v/v) fetal bovine serum in PBST. To visualize subcellular localization of exogenous MTCL2, cells were treated with modified PBST containing 0.5% TritonX-100 instead of Tween 20 for 10 min after methanol fixation. To examine different fixation conditions, cells were fixed with 4% paraformaldehyde in BRB80 buffer (80 mM PIPES-KOH [pH 6.8], 1 mM MgCl_2_, 1 mM EGTA), with or without pre-extraction using BRB80 buffer supplemented with 4 mM EGTA and 0.5% TX-100, for 30 s at 37 °C. After fixation and blocking, samples were incubated with appropriate primary antibodies diluted in TBST (10 mM Tris–HCl [pH 7.5], 150 mM NaCl, 0.01% [v/v] Tween 20) containing 0.1% (w/v) BSA for 45 min at RT, except for MTCL1, MTCL2, and MAP4 staining, which was performed overnight at 4 °C. After washing with PBST, samples were visualized with the appropriate secondary antibodies conjugated with Alexa Fluor 488, 555, or 647 (Life Technologies Corporation) by incubating for 45 min at RT. Antibodies were diluted as follows: anti-KIAA0802 (1/1000), anti-SOGA1 (1/2000), anti-α-tubulin (1:1000), anti-β-tubulin (1:2000), anti-acetylated tubulin (1:1000), anti-V5 (1:4000), anti-GM130 (1:1000), anti-GS28 (1:300), anti-GFP (1:2,000), anti-MAP4 (1:1000), anti-pericentrin (1:1000), anti-CLASP1 (1:500), anti-CLASP2 (1:500), anti-giantin (1:1000), anti-GCC185 (1:2000; anti-AKAP450 (1:500); anti-Golgin97 (1:1000). All secondary antibodies were used at a 1:2000 dilution. The nuclei were counterstained with 4’, 6-diamidino-2-phenylindole (MBL, Japan) at a 1:2000 dilution in PBST during the final wash. For image acquisition, samples on coverslips were mounted onto glass slides in Prolong Diamond Antifade Mountant (Thermo Fisher Scientific).

### Image acquisition and processing

High-resolution images were acquired using a Leica SP8 laser scanning confocal microscopy system equipped with an HC PL APO 63x/1.40 Oil 2 objective, using the Hybrid Detector in photon-counting mode. To obtain super-resolution images, HyVolution2 imaging was performed on the same system using the Huygens Essential software (Scientific Volume Imaging) (Borlinghaus & Kappel, 2016). To obtain wide-view images for quantification (Fig. 5, Figs. EV4 and 5), conventional fluorescence images were obtained using an AxioImager ZI microscope (Carl Zeiss, Oberkochen, Germany) equipped with a Plan APCHROMAT 40×/0.95 objective using an Orca II CCD camera (Hamamatsu Photonics, Shizuoka, Japan). The laterally expanding angle of the Golgi apparatus around the nuclei and the skewness of pixel intensity were quantified using the “Measure” function of ImageJ software. For statistical analysis, photographs of several fields containing ∼40 cells with similar densities were taken. All cells in each field were subjected to the following quantification analysis to avoid selection bias. In rescue experiments, ∼100 cells expressing exogenous MTCL2 at similar expression levels as the endogenous one were collected from ∼10 fields with similar cell densities. For live-cell imaging, differential interference contrast images were acquired using a Leica SP8 confocal microscopy system equipped with an HCX PL APO 10×/0.40 objective using a 488 nm laser line. Areas newly covered by migrated cells during wound healing for 440 min were estimated using the “Measure” function of ImageJ software and normalized by the length of the corresponding wound edge at time 0.

### MT-binding assay

MBP or MBP-mMTCL1 CT1 was purified from the soluble fraction of *E. coli* according to the standard protocol and dialyzed against BRB80 buffer. Each MBP was incubated with taxol-stabilized MTs (final concentrations of both the sample protein and α/β-tubulin heterodimer were 0.5 mg/mL) in BRB80 supplemented with 1.5 mM MgCl_2_ and 1 mM GTP for 15 min at RT and subjected to centrifugation (200,000 × *g*) for 20 min at 25°C on a cushion of 40% glycerol in BRB buffer. Following careful removal of the supernatant and glycerol cushion, the resultant MT pellet was gently washed with PBST three times and solubilized with SDS sample buffer (10% β-mercaptoethanol, 125 mM Tris-HCl [pH 6.8], 2% SDS, 10% glycerol, and 0.005% bromophenol blue) for subsequent SDS-PAGE analysis.

### Pull-down experiments

HEK293T cells (∼8 × 10^6^ cells) transfected with appropriate expression vectors were solubilized in 500 μL lysis buffer (20 mM Tris-HCl [pH 7.5], 150 mM NaCl, 0.3% TX-100, 2 mM MgCl_2_, 1 mM EGTA) containing a cocktail of protease and phosphatase inhibitors (Roche Applied Science) for 30 min at 4°C. They were then briefly sonicated and centrifuged at 15,000 × *g* for 30 min. The resulting supernatants were mixed with streptavidin-conjugated magnetic beads (Cytiva) for ∼2 h at 4°C. The beads were collected using a magnet, washed with lysis buffer three times, and then boiled in SDS sample buffer. Proteins released from the beads were subjected to western blotting analysis using the following antibodies: anti-V5 (1:3000), anti-GFP (1:1000), and anti-Flag (1:3000).

## Acknowledgments

The authors thank Yoshihiro Miwa (University of Tsukuba) and Ikuko Hayashi (Yokohama City University) for providing the pOSTet15.1 expression vector and GFP-CLASP2α expression vector, respectively. This work was supported by KAKENHI (to A.S.) (Grant Numbers 16H04765 and 19H03228) of the Ministry of Education, Culture, Sports, Science, and Technology (MEXT) of Japan.

## Author contributions

A.S. planned and performed the experiments, interpreted the results, and wrote the manuscript. R.M., M. M., S. M., C. Y., and Y.I. performed the experiments.

## Expanded View Figure Legends

**Figure EV1.**
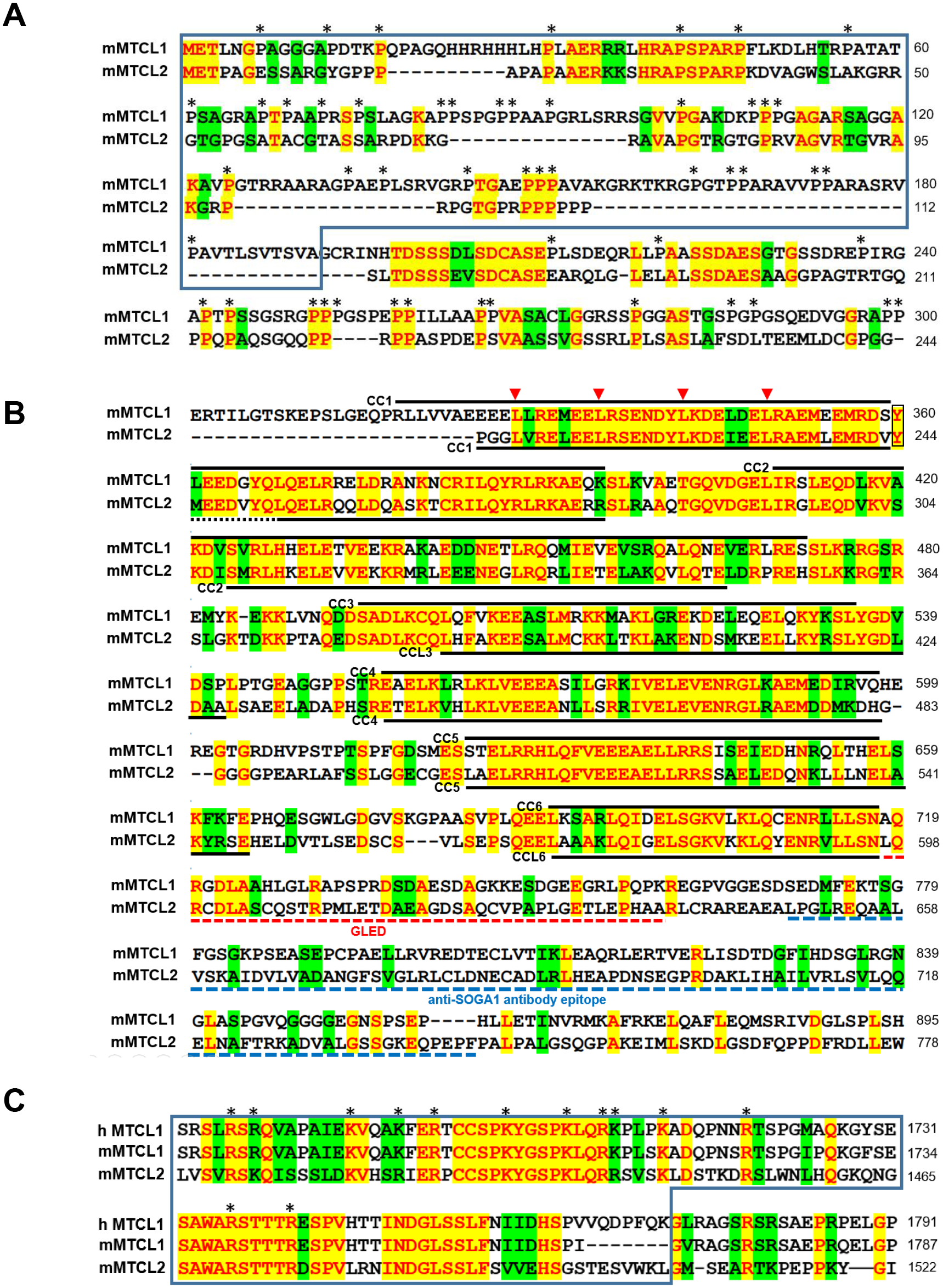
Sequence alignment of amino acid sequences of mouse MTCL1 and 2. A. The N-terminal sequences. Boxed region corresponds to N-MTBD of MTCL1. Asterisks indicate the positions of proline highly condensed in this region. B. The N-terminal coiled-coil region. The positions of each coiled-coil motif (CC) or coiled-coil-like motif (CCL) of MTCL1 or 2 are indicated by bold lines on the top or bottom of each sequence, respectively. GLED sequence of MTCL2 is underlined by a red dashed line. Four leucine residues mutated in 4LA or 4LP mutants are indicated red arrowheads. A tyrosine residue that disrupts the periodicity of CC1 is boxed. Blue dotted lines indicate the region corresponding to the epitope for anti-SOGA1 antibody C. The sequences of the C-terminal MT-binding regions. Because MTCL1 C-MTBD (boxed) was defined for human protein (Sato *et al*, 2013), the human sequence of MTCL1 is also included in this alignment. The region of mouse MTCL2 corresponding to MTCL1 C-MTBD is designated the “KR-rich region” since the conserved basic residues (asterisks) are condensed.

**Figure EV2.**
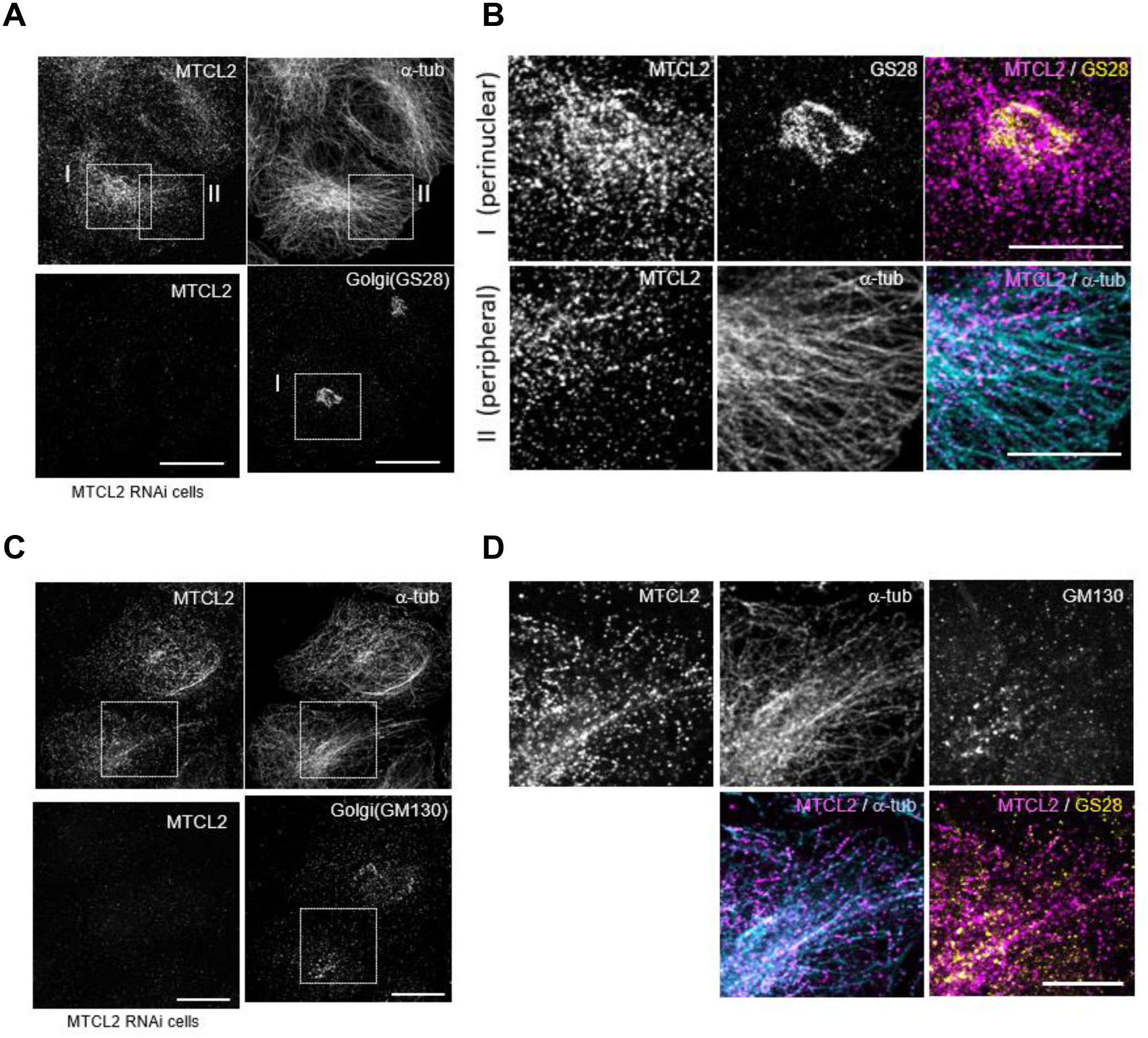
Fixation conditions did not affect the staining pattern of MTCL2. A. HeLa-K cells fixed with 4% paraformaldehyde were stained with anti-SOGA1 (MTCL2) together with anti-α-tubulin and anti-GS28 antibody. The specificity of anti-SOGA1 signals is indicated by their disappearance in MTCL*2*-knockdown cells subjected to the same procedures (see a lower left panel). Scale bar: 20 μm. B. Boxed regions in (A) are enlarged to examine the colocalization of MTCL2 on the GA and MTs more closely. Scale bar: 10 μm. C. HeLa-K cells were fixed with 4% paraformaldehyde after brief treatment of an extraction buffer containing 0.5% TX-100 and 4 mM EGTA. The specificity of anti-SOGA1 staining signals is indicated by their disappearance in *MTCL2*-knockdown cells subjected to the same procedures (see a lower left panel). Scale bar: 20 μm. D. The boxed region in (C) is enlarged to examine the colocalization of MTCL2 on the Golgi and MTs more closely. Scale bar: 10 μm.

**Figure EV3.**
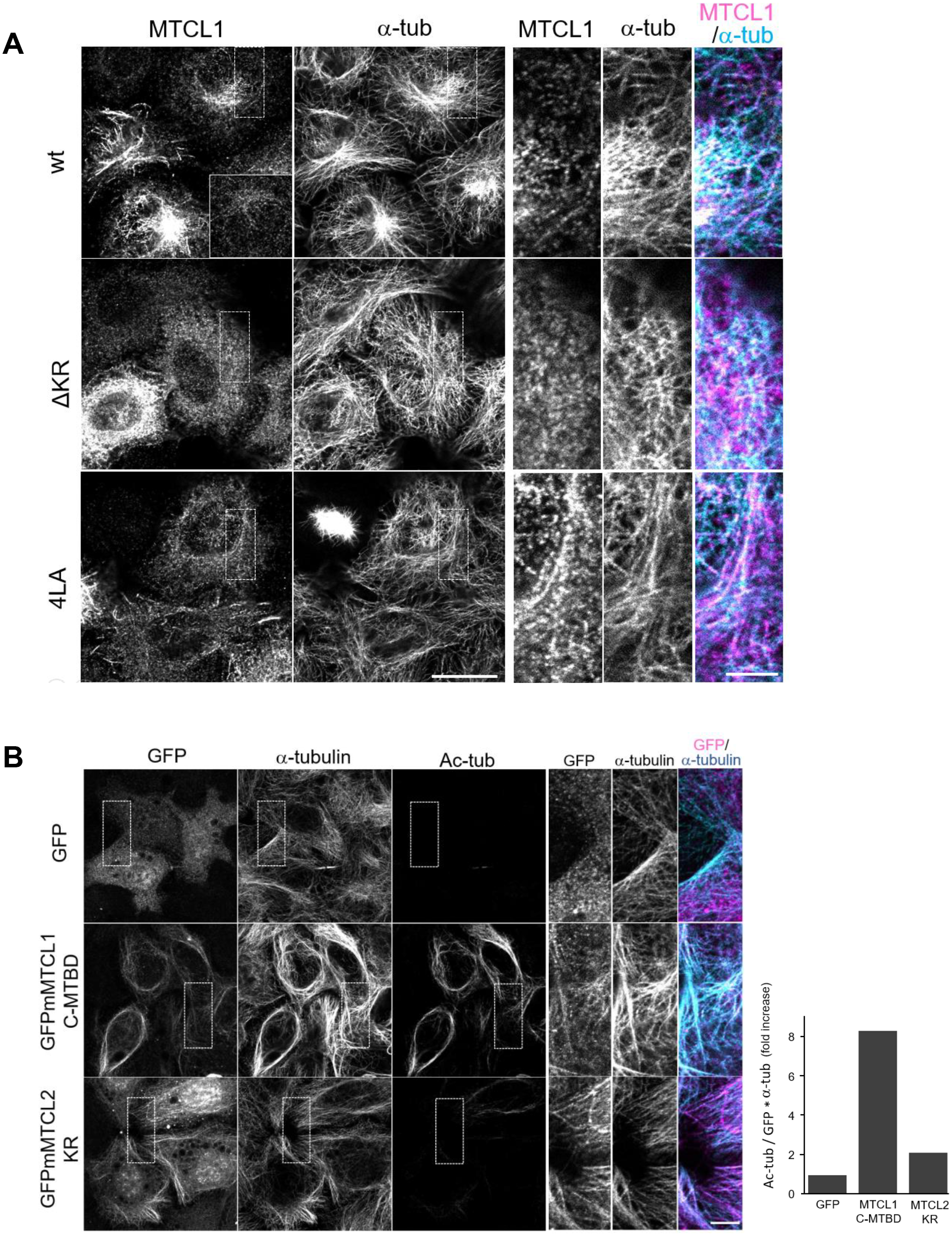
KR-rich region is the MT-binding region of MTCL2. A. Subcellular localization of exogenously expressed mMTCL2 and its mutants in MTCL2-knockdown HeLa-K cells. Scale bar: 20 μm. The cells expressing exogenous MTCL2 at a level several-fold higher than the endogenous MTCL2 level are shown (see an inset in an upper left panel in which a staining image of endogenous MTCL2 in a HeLa-K cell is shown under the same condition). Boxed regions are enlarged in right panels. Scale bar: 5 μm. Note that MTCL2 ΔKR but not 4LA mutant lost MT association activities. Intriguingly, in contrast to the N fragment (Fig. 3), ΔKR mutant had no Golgi localization, suggesting that MT binding through the KR-rich region was a prerequisite for the association of full-length MTCL2 with the Golgi membrane. B. HeLa-K cells exogenously expressing GFP, GFP-mMTCL1 C-MTBD, or GFP-mMTCL2 KR were stained with the indicated antibodies. Scale bar: 20 μm. Boxed regions are enlarged in right panels. Scale bar: 5 μm. Note that, in contrast to MTCL2 KR, MTCL1 C-MTBD strongly induced tubulin acetylation and MT bundling. Quantitative comparison of acetylated tubulin signals in the insets after normalization by GFP and α-tubulin signal intensities is shown on the left.

**Figure EV4.**
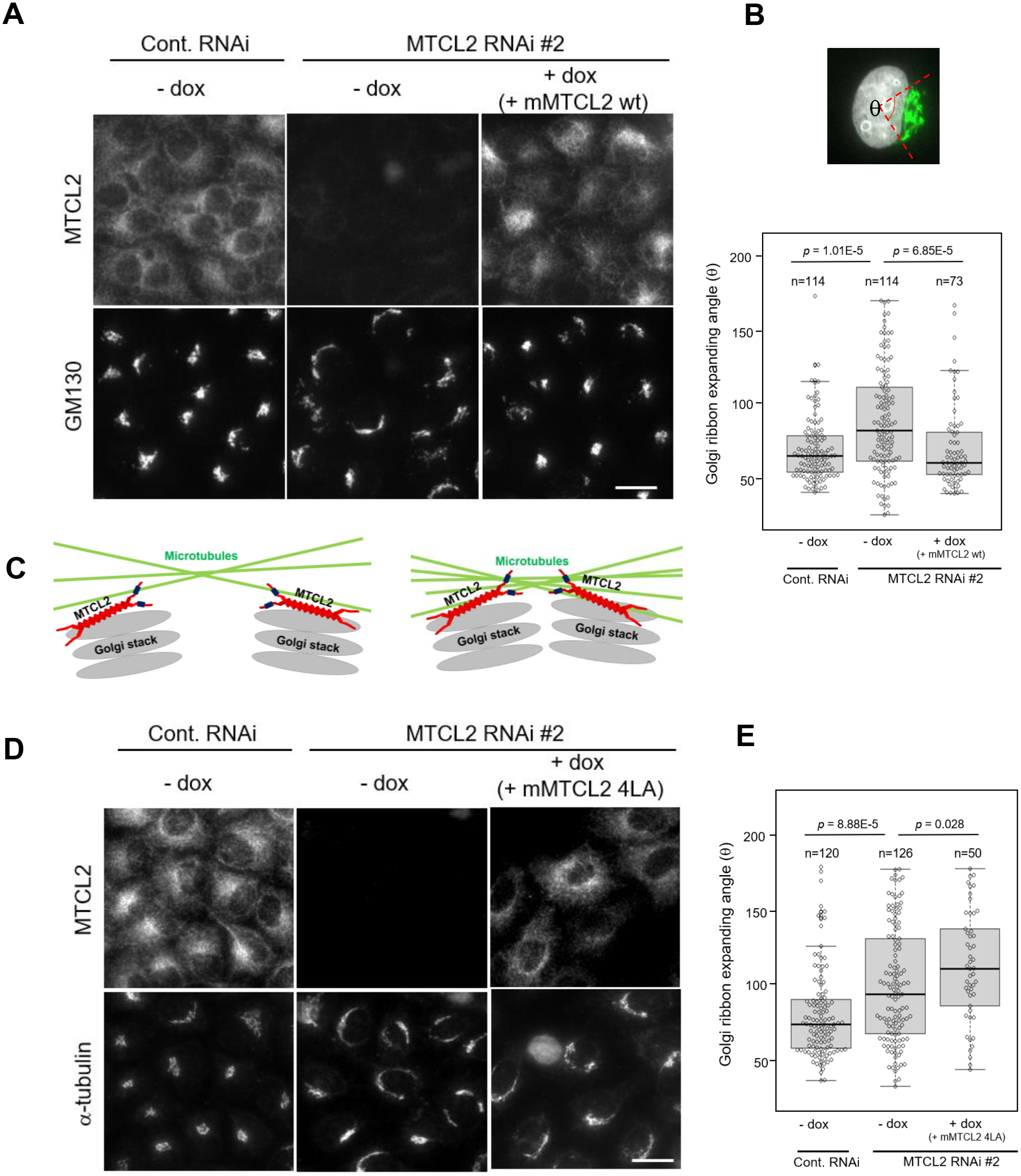
MTCL2 promotes clustering of the Golgi stacks in a Golgi-association-dependent manner. A. HeLa-K cells stably harboring pOSTet15.1 expression vector for mouse MTCL2 were transfected with siRNAs for control or MTCL2 knockdown (#2) in the presence or absence of 100 nM doxycycline and doubly stained with anti-SOGA1 (MTCL2) and anti-GM130 antibodies, as indicated on the left. Note that cells subjected to control RNAi show compact Golgi ribbon structures at one side of the perinuclear region. Such Golgi ribbon structures become laterally expanded around the nucleus in MTCL2-knockdown cells (-dox), whereas exogenous expression of RNAi-resistant MTCL2 (+dox) strongly restores their compactness. Scale bar: 20 μm. B. Quantification of Golgi ribbon expanding angle (θ) around the nuclei (top panel) in each condition. Bottom is a box plot of the angle distribution in each condition. The lines within each box represent medians. Data represent the results of the indicated number (n) of cells from a typical experiment (biological replicates). The *p* values were estimated using the Wilcoxon test. Statistical data of technical replicates (three independent experiments) are demonstrated in Appendix Fig. S4. C. A model explaining how MT accumulation secondarily increases clustering of individual Golgi stacks. D. HeLa-K cells stably harboring pOSTet15.1 expression vector for mouse MTCL2 4LA were subjected to the same experimental procedure as in (A). Note that compactness of Golgi ribbon was not restored by expression of mouse MTCL2 4LA. Scale bar: 20 μm. E. Quantitative analysis as in (B).

**Figure EV5.**
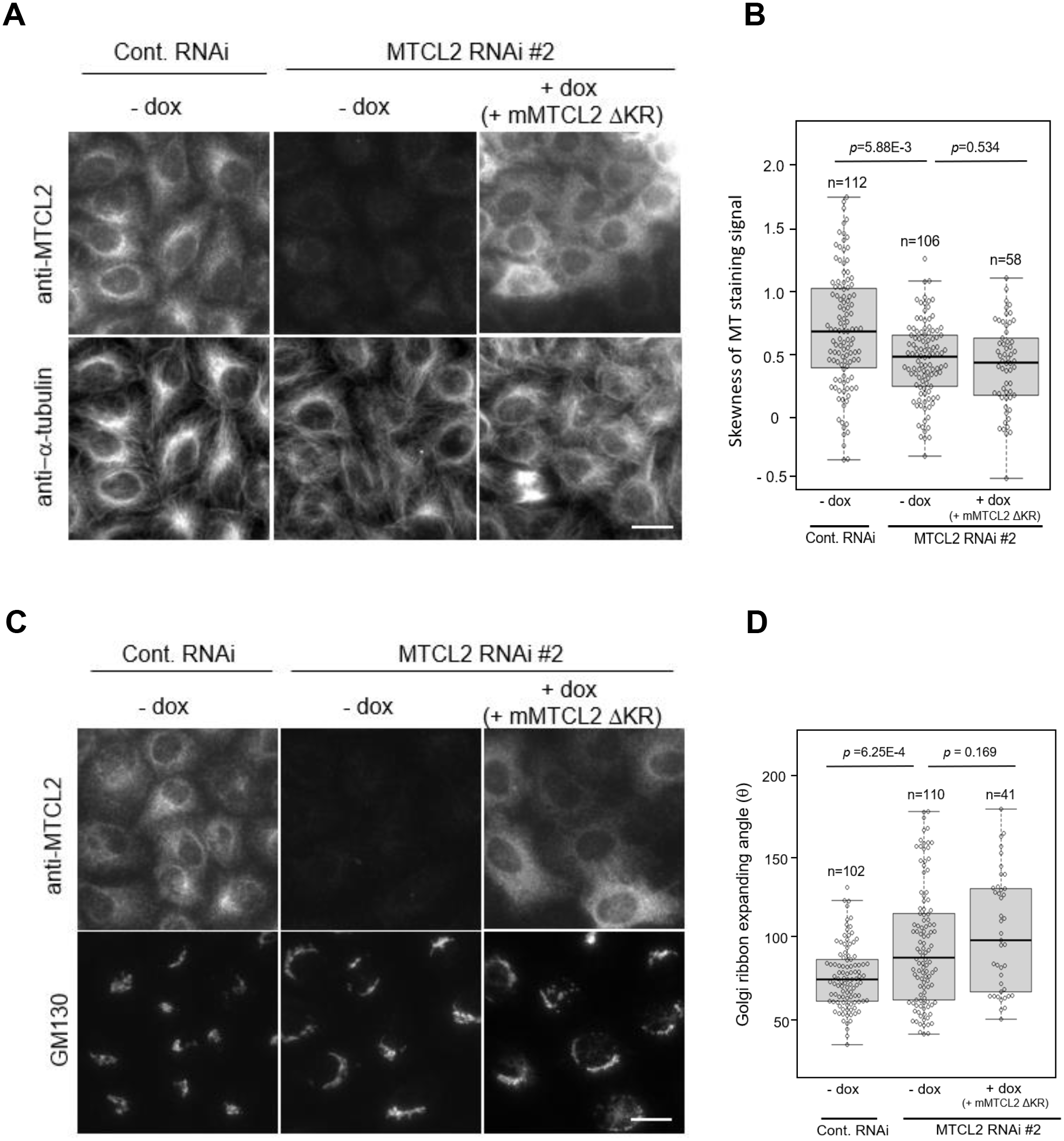
MT-binding activity is also required for MTCL2 function to facilitate the perinuclear accumulation of MTs and the Golgi ribbon compactness. HeLa-K cells stably harboring pOSTet15.1 expression vector for mouse MTCL2 ΔKR were transfected with siRNAs for control or MTCL2 knockdown (#2) in the presence or absence of 100 nM doxycycline (dox). Perinuclear accumulation of MTs (A and B) and expansion of the Golgi ribbon around the nucleus (C and D) were analyzed in the same manner as described in Fig. 5 and Fig. EV4 legends. Scale bar: 20 μm.

## Expanded View Movie Legends

**Movie EV1. Wound healing of RPE1 cells subjected to control knockdown.** Differential interference contrast images of cells were taken every 10 min for 440 min. The video speed is 6 fps. Representative frames of this movie are shown in Fig. 6B.

**Movie EV2. Wound healing of RPE1 cells subjected to *MTCL2* knockdown.** Data were collected as described in the Supplementary Movie 1 legend. Representative frames of this movie are shown in Fig. 6B.

**Appendix Figure S1.**
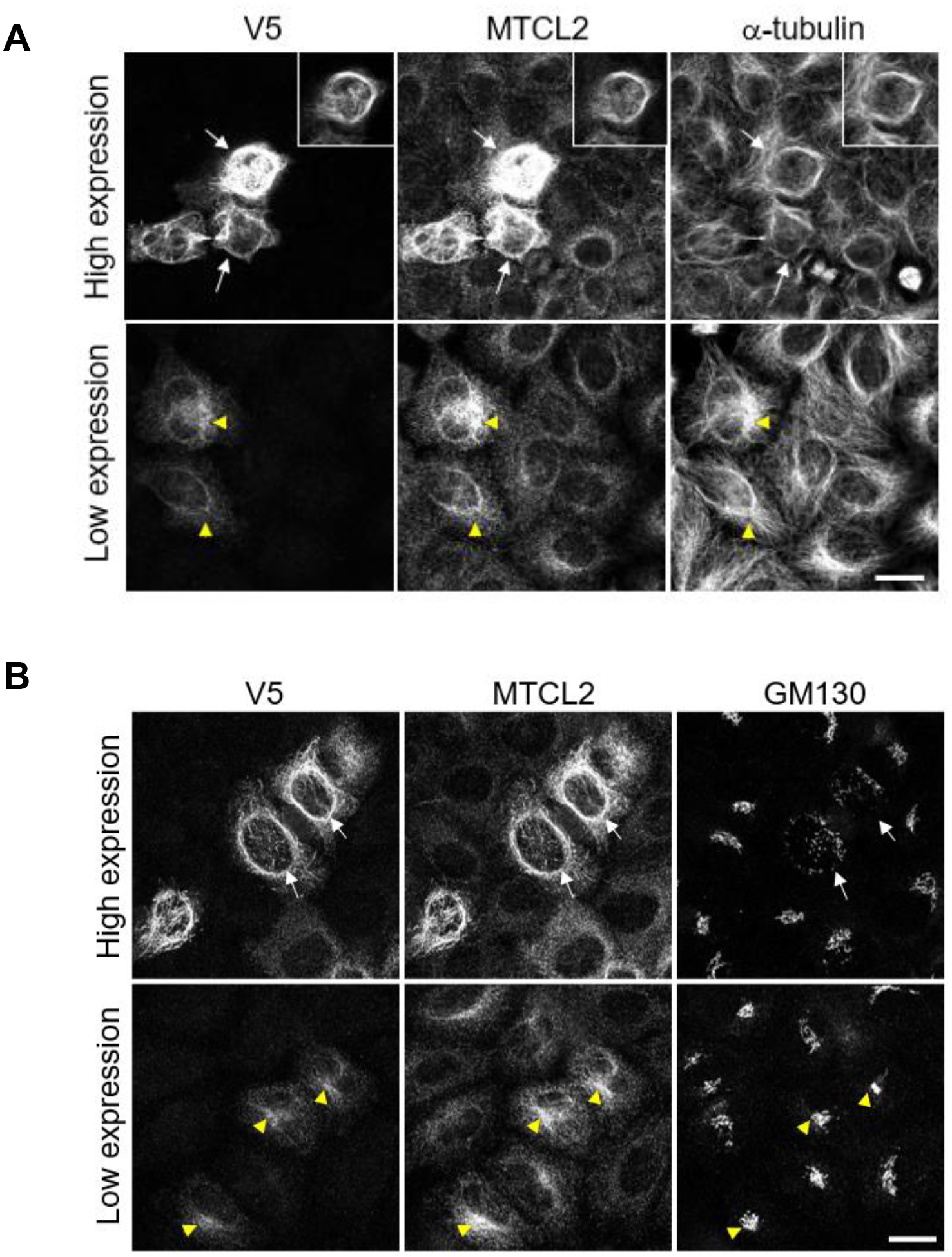
Localization of exogenously expressed MTCL2 mimics that of endogenous proteins at low expression levels. HeLa-K cells stably harboring 6xV5-tagged mouse MTCL2 expression vector (pOSTet15.1) were cultured in the presence of 100 ng/mL doxycycline and stained with the indicated antibodies. Scale bar: 20 μm. Arrows indicate cells highly expressing exogenous MTCL2, whereas yellow arrowheads indicate cells expressing exogenous MTCL2 at a level comparable to endogenous MTCL2. The insets in (A) show alternative images of a cell located at the center of the panel, in which contrasts of the individual staining signals are adjusted separately to provide unsaturated images.

**Appendix Figure S2.**
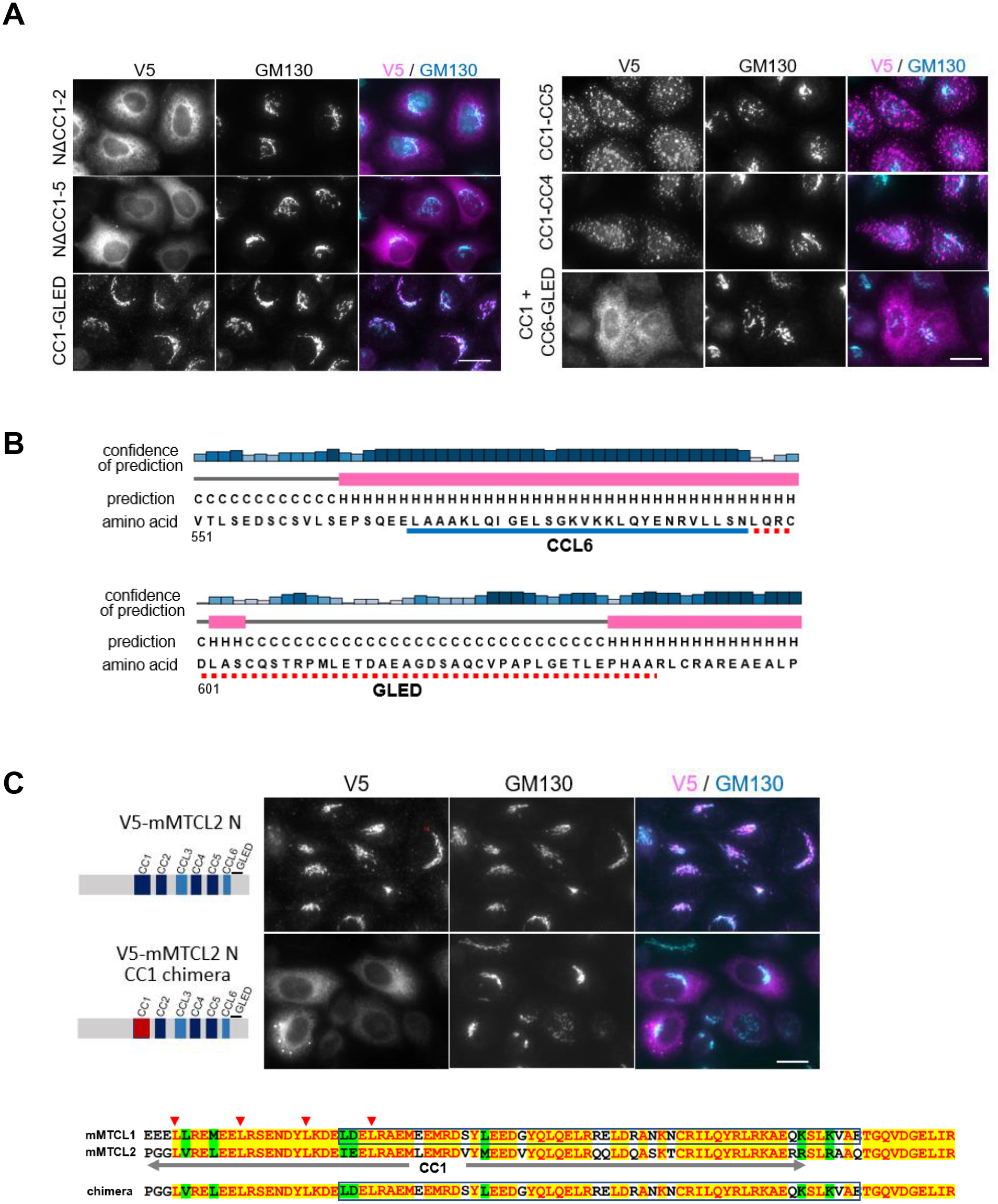
The essential sequence required for the Golgi association of the MTCL2 N fragment. A. Subcellular localization of the indicated mutants expressed in HeLa-K cells ( Fig. 4A). Scale bar: 20 μm. B. The amino acid sequence of GLED and its secondary structure redicted using PSIPED (http://bioinf.cs.ucl.ac.uk/psipred/). C. Subcellular localization of the CC1 chimera of the N fragment, in which the highly conserved CC1 sequence of MTCL2 was seamlessly exchanged with that of MTCL1. Scale bar, 20 μm. The amino acid sequence of CC1 in the chimera mutant is shown below.

**Appendix Figure S3.**
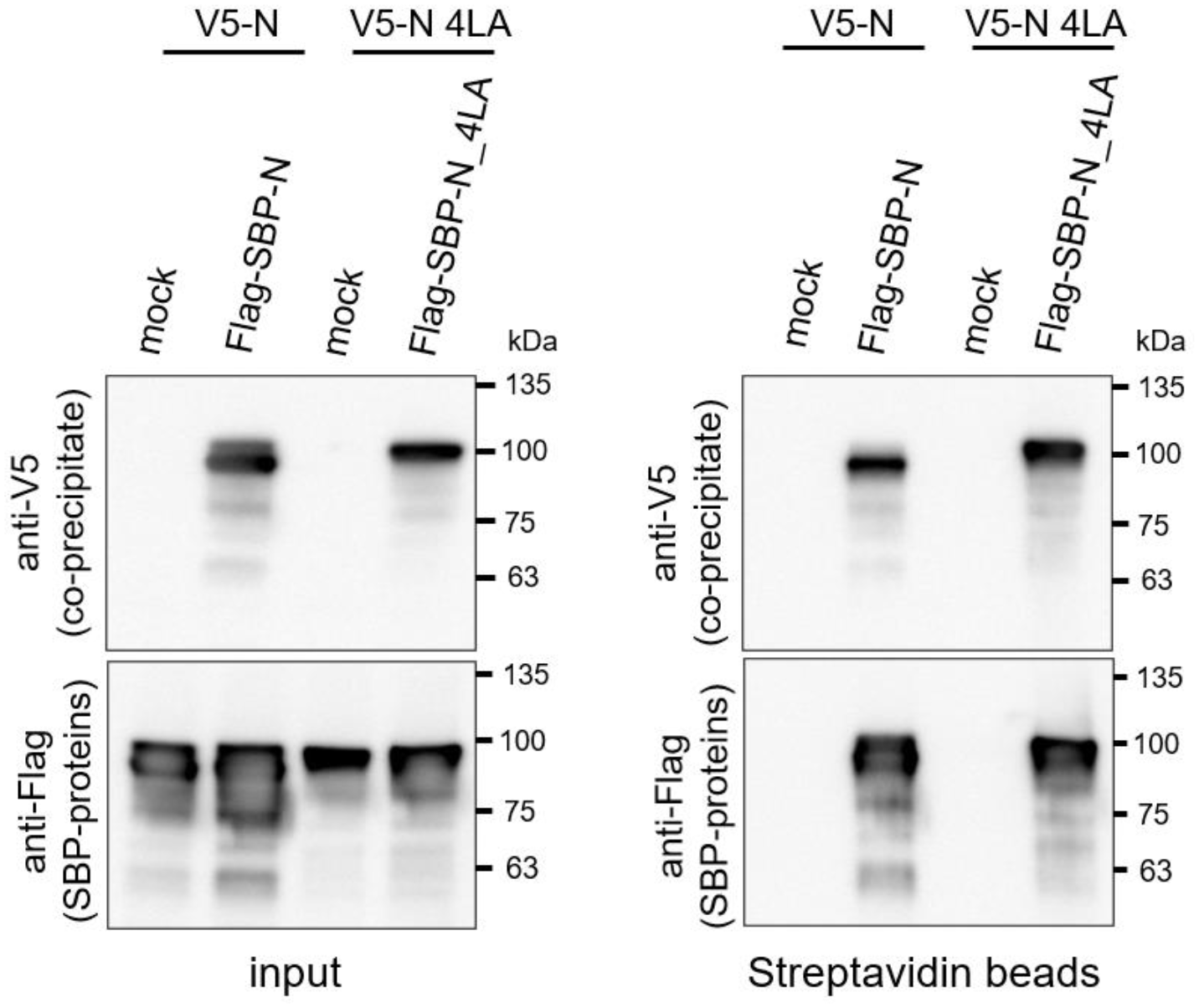
The 4LA mutation does not interfere with the homo-oligomerization activity of N fragment. A streptavidin pull-down experiment was performed for soluble extracts (input) of HEK293 cells expressing V5-N with Flag-SBP-N or V5-N 4LA with Flag-SBP-N 4LA, as indicated. In mock samples, empty backbone vectors for Flag-SBP constructs were transfected with each V5 construct.

**Appendix Figure S4.**
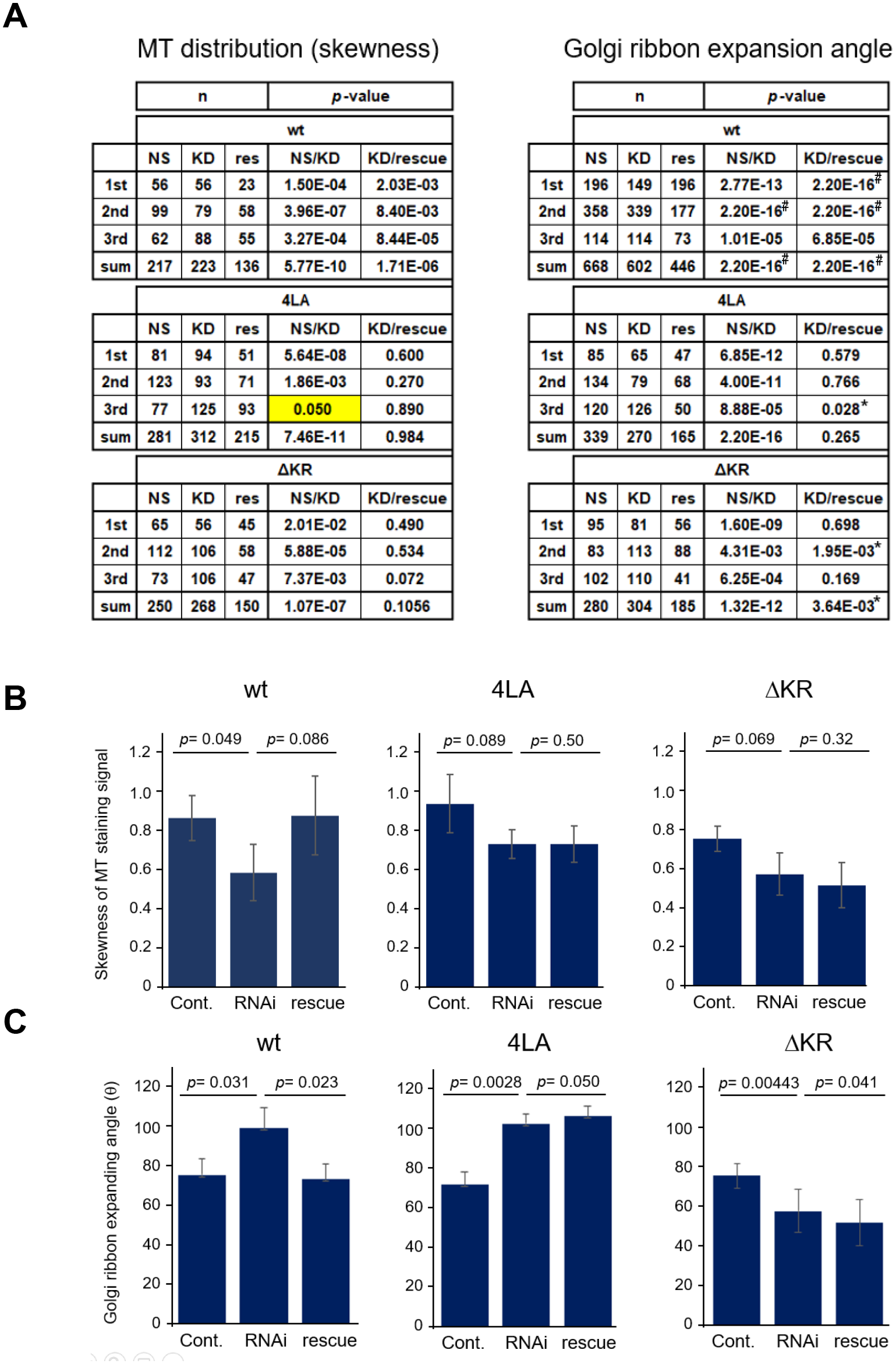
Statistical data for technical replicates of the rescue experiments. A. Numbers of biological replicates (n) and *p* values estimated by the Wilcoxon test are listed for each rescue experiment replicated three times. Left, experiments used to examine rescue activity for MT distribution. Right, Golgi ribbon compactness. The *p* values indicated by # mean less than 2.20e-16. Expression of MTCL2 mutants (4LA, ΔKR) tended to worsen the knockdown phenotypes of MTCL2, sometimes resulting in low *p* values in KD/rescue comparison, as indicated by asterisks. Note that essential trends of each MTCL2 mutant shown in Fig. 5 and Figs. EV4 and 5 are highly reproduced except in an experiment (yellow cell) in which the MTCL2-knockdown effect was rather low. B, C. Mean of biological replicates in each experiment listed in (A) was averaged in three technical replicates and compared between each condition. Data represent the mean ± S.D. of three independent experiments for MT distribution (B) and Golgi ribbon compactness (C). The *p* value was estimated using Student’s t-test assuming a one-tailed distribution and two-sample unequal variance.

**Appendix Figure S5.**
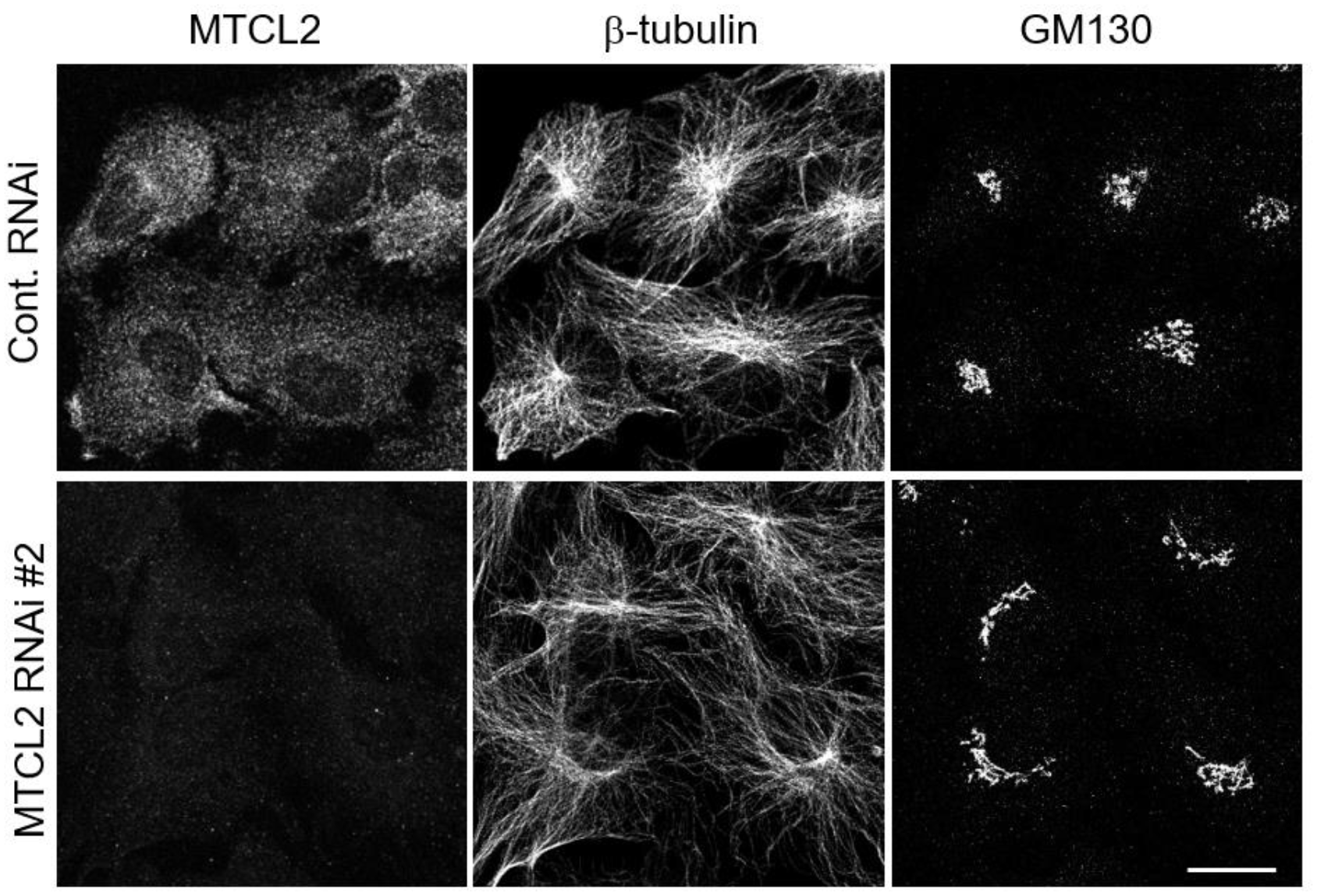
Effects of *MTCL2* knockdown on RPE1 cells. RPE1 cells transfected with control or MTCL2 siRNAs were subjected to immunofluorescence analysis using the indicated antibodies. Note that reduced accumulation of MTs around the Golgi and lateral expansion of the Golgi ribbon were observed in this cell line. Scale bar: 20 μm.

**Appendix Figure S6.**
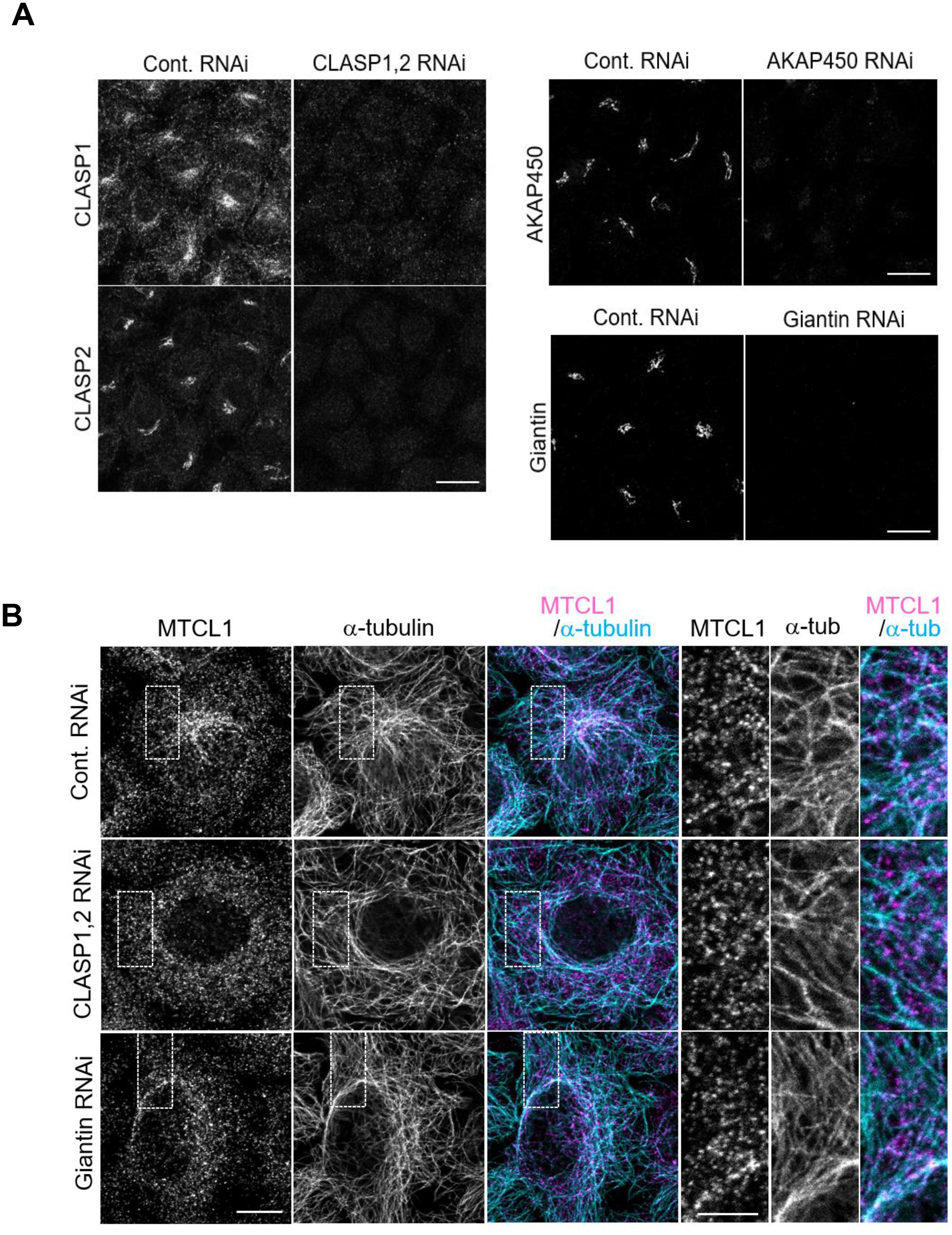
Effects of CLASP, AKAP450, and giantin knockdown through RNAi in cells. A. Reduced expression of target proteins of the indicated siRNAs is shown. Scale bar: 20 μm. B. Colocalization of endogenous MTCL2 with MTs in the indicated knockdown cells was examined in HeLa-K cells. Scale bar: 10 μm. Boxed regions are enlarged in the right panels. Scale bar: 5 μm.

**Appendix Figure S7.**
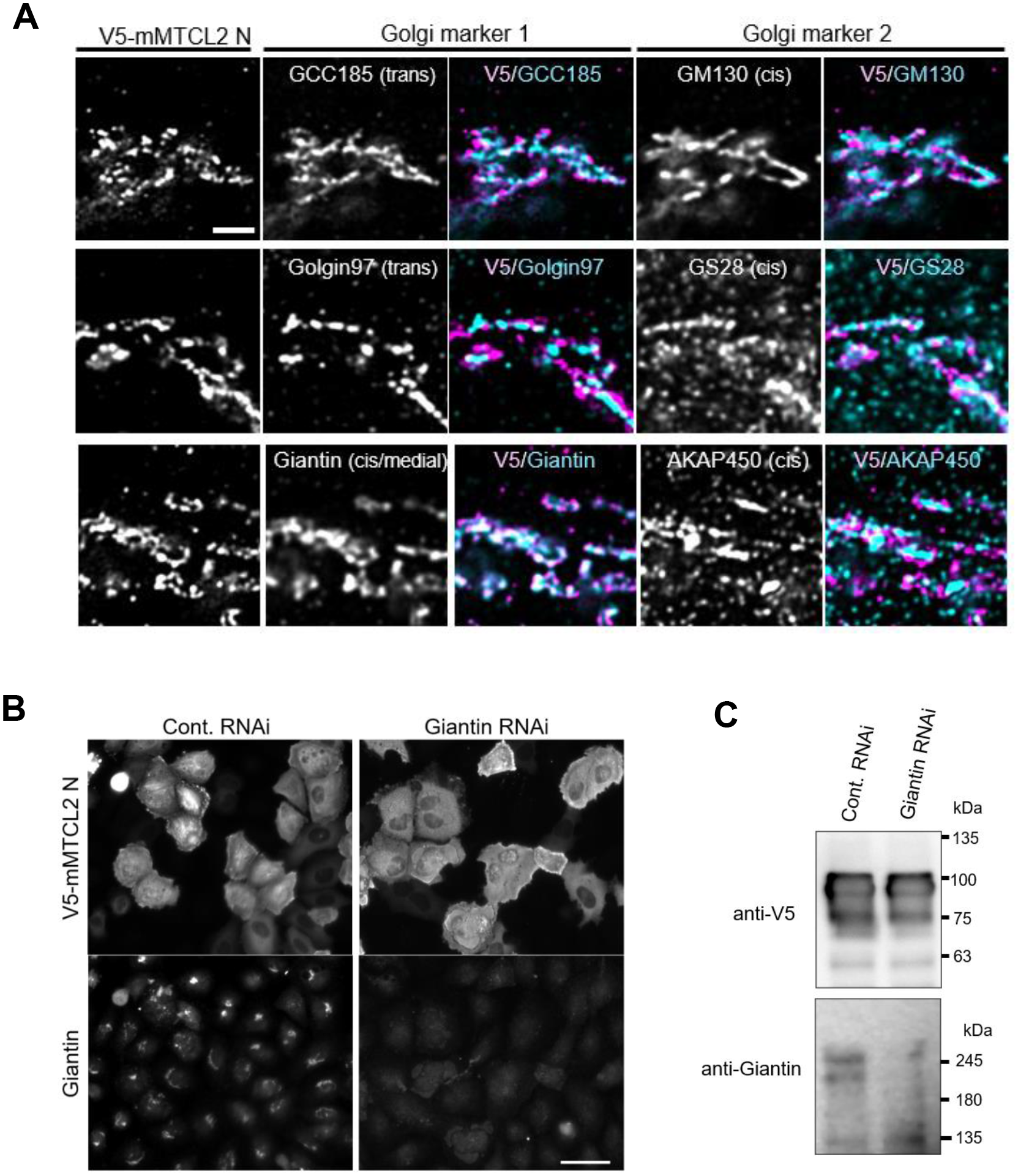
Giantin is involved in the Golgi association of the MTCL2 N fragment. A. Subcellular localization of V5-mMTCL2 N fragment in HeLa-K cells was compared with that of Golgi-resident proteins using super-resolution microscopy. Scale bar: 2μm. Note that the N-terminal fragment of MTCL2 shows colocalization with the cis/medial Golgi protein giantin/GOLGB1 most clearly. The fragment showed distinct localization from cis Golgi marker proteins, suggesting that it is mainly associated with the medial Golgi cisternae. B. Levels of V5-mMTCL2 N fragment in control and giantin-knockdown cells were compared through immunostaining analysis using the indicated antibodies after paraformaldehyde fixation, which prevented leakage of cytosolic protein during fixation. Scale bar: 50 μm. C. Levels of V5-mMTCL2 N fragment in control and giantin-knockdown cells were compared through western blotting analysis using total cell extracts.

**Appendix Figure S8.**
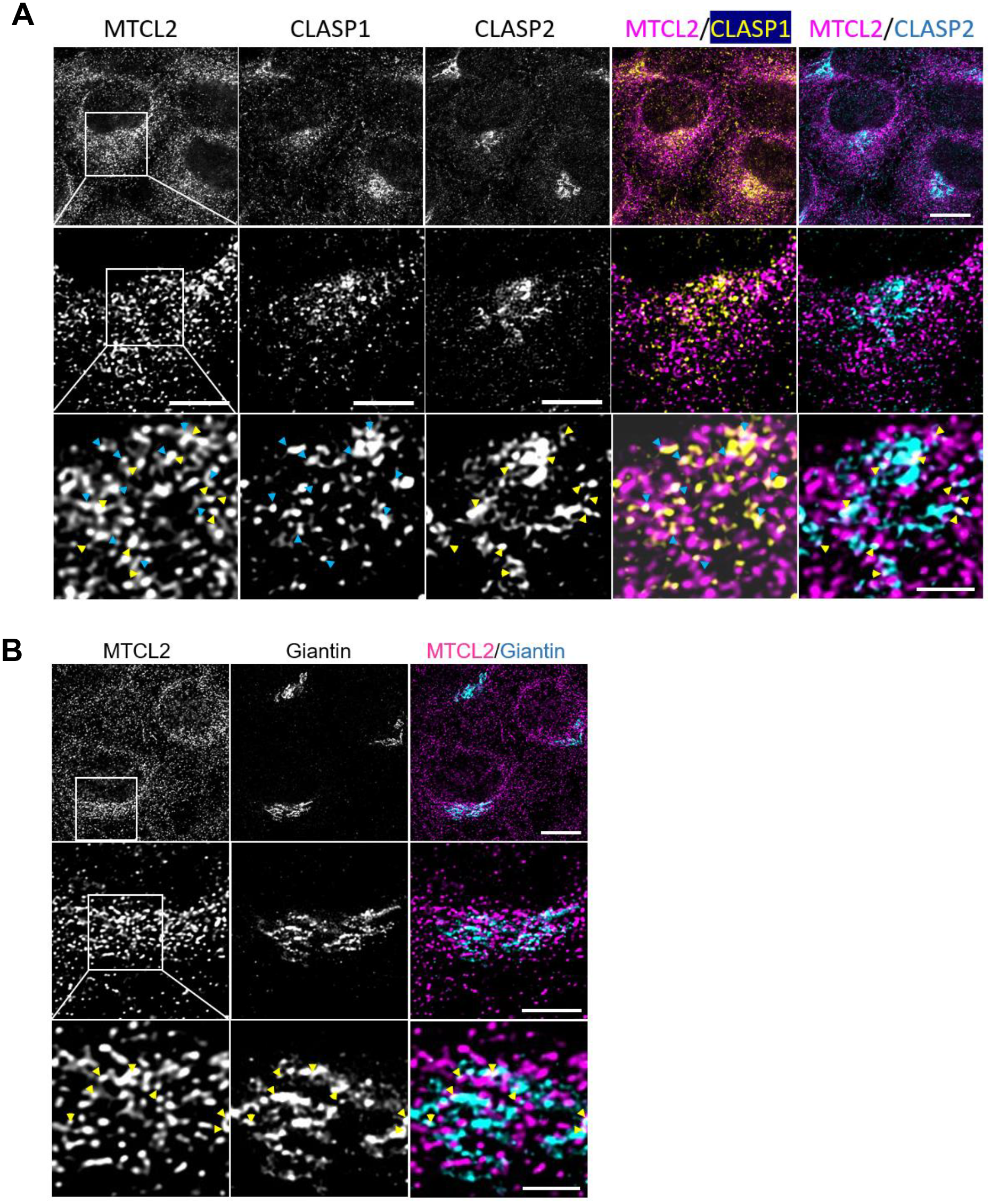
Endogenous MTCL2 exhibited partial colocalization with CLASPs and giantin. Subcellular localization of endogenous MTCL2 in HeLa-K cells was compared with that of CLASPs (A) and giantin (B) using super-resolution microscopy. Boxed regions are serially enlarged in the middle and bottom panels. Arrowheads indicate the regions where each protein shows colocalization with MTCL2. Scale bars: 10 μm (top), 5 μm (middle), and 2 μm (bottom).

## References

Abdul Kader M, Satake T, Yoshida M, Hayashi I & Suzuki A (2017) Molecular basis of the microtubule-regulating activity of microtubule crosslinking factor 1. PLoS One 12: e0182641

Alvarez C, Garcia-Mata R, Hauri HP & Sztul E (2001) The p115-interactive Proteins GM130 and Giantin Participate in Endoplasmic Reticulum-Golgi Traffic. J Biol Chem 276: 2693–2700

Bartolini F & Gundersen GG (2006) Generation of noncentrosomal microtubule arrays. J Cell Sci 119: 4155–63

Bergmann JE, Kupfer A & Singer SJ (1983) Membrane insertion at the leading edge of motile fibroblasts. Proc Natl Acad Sci U S A 80: 1367–1371

Borlinghaus RT & Kappel C (2016) HyVolution—the smart path to confocal super-resolution. Nat Methods 13: i–iii

Chabin-Brion K, Marceiller J, Perez F, Settegrana C, Drechou A, Durand G & Poüs C (2001) The Golgi complex is a microtubule-organizing organelle. Mol Biol Cell 12: 2047–60

Chapin SJ & Bulinski JC (1991) Non-neuronal 210 x 10(3) Mr microtubule-associated protein (MAP4) contains a domain homologous to the microtubule-binding domains of neuronal MAP2 and tau. J Cell Sci 98 ( Pt 1): 27–36

Combs TP & Marliss EB (2014) Adiponectin signaling in the liver. Rev Endocr Metab Disord 15: 137–147

Conduit PT, Wainman A & Raff JW (2015) Centrosome function and assembly in animal cells. Nat Rev Mol Cell Biol 16: 611–624

Cowherd RB, Asmar MM, Alderman JM, Alderman E a, Garland AL, Busby WH, Bodnar WM, Rusyn I, Medoff BD, Tisch R, et al (2010) Adiponectin lowers glucose production by increasing SOGA. Am J Pathol 177: 1936–1945

Efimov A, Kharitonov A, Efimova N, Loncarek J, Miller PM, Andreyeva N, Gleeson P, Galjart N, Maia ARR, McLeod IX, et al (2007) Asymmetric CLASP-Dependent Nucleation of Noncentrosomal Microtubules at the trans-Golgi Network. Dev Cell 12: 917–930

Hurtado L, Caballero C, Gavilan MP, Cardenas J, Bornens M & Rios RM (2011) Disconnecting the Golgi ribbon from the centrosome prevents directional cell migration and ciliogenesis. J Cell Biol 193: 917–33

Kruse R, Krantz J, Barker N, Coletta RL, Rafikov R, Luo M, Højlund K, Mandarino LJ & Langlais PR (2017) Characterization of the CLASP2 protein interaction network identifies SOGA1 as a microtubule-associated protein. Mol Cell Proteomics 16: 1718–1735

Linstedt AD, Foguet M, Renz M, Seelig HP, Glick BS & Hauri HP (1995) A C-terminally-anchored Golgi protein is inserted into the endoplasmic reticulum and then transported to the Golgi apparatus. Proc Natl Acad Sci U S A 92: 5102–5105

Meiring JCM, Shneyer BI & Akhmanova A (2020) Generation and regulation of microtubule network asymmetry to drive cell polarity. Curr Opin Cell Biol 62: 86– 95

Miller PM, Folkmann AW, Maia ARR, Efimova N, Efimov A & Kaverina I (2009) Golgi-derived CLASP-dependent microtubules control Golgi organization and polarized trafficking in motile cells. Nat Cell Biol 11: 1069–80

Mimori-Kiyosue Y, Grigoriev I, Lansbergen G, Sasaki H, Matsui C, Severin F, Galjart N, Grosveld F, Vorobjev I, Tsukita S, et al (2005) CLASP1 and CLASP2 bind to EB1 and regulate microtubule plus-end dynamics at the cell cortex. J Cell Biol 168: 141–153

Nishita M, Satake T, Minami Y & Suzuki A (2017) Regulatory mechanisms and cellular functions of non-centrosomal microtubules. J Biochem 109: 20029–20034

Rivero S, Cardenas J, Bornens M & Rios RM (2009) Microtubule nucleation at the cis-side of the Golgi apparatus requires AKAP450 and GM130. EMBO J 28: 1016– 1028

Sanders AAWM & Kaverina I (2015) Nucleation and Dynamics of Golgi-derived Microtubules. 9: 1–7

Satake T, Yamashita K, Hayashi K, Miyatake S, Tamura-Nakano M, Doi H, Furuta Y, Shioi G, Miura E, Takeo YH, et al (2017) MTCL1 plays an essential role in maintaining Purkinje neuron axon initial segment. EMBO J 36: 1227–1242

Sato Y, Akitsu M, Amano Y, Yamashita K, Ide M, Shimada K, Yamashita A, Hirano H, Arakawa N, Maki T, et al (2013) The novel PAR-1-binding protein MTCL1 has crucial roles in organizing microtubules in polarizing epithelial cells. J Cell Sci 126: 4671–4683

Sato Y, Hayashi K, Amano Y, Takahashi M, Yonemura S, Hayashi I, Hirose H, Ohno S & Suzuki A (2014) MTCL1 crosslinks and stabilizes non-centrosomal microtubules on the Golgi membrane. Nat Commun 5: 5266

Sönnichsen B, Lowe M, Levine T, Jämsä E, Dirac-Svejstrup B & Warren G (1998) A role for giantin in docking COPI vesicles to Golgi membranes. J Cell Biol 140: 1013–1021

Tanaka J, Miwa Y, Miyoshi K, Ueno A & Inoue H (1999) Construction of Epstein-Barr virus-based expression vector containing mini-oriP. Biochem Biophys Res Commun 264: 938–43

Vorobjev IA & Nadezhdina ES (1987) The Centrosome and Its Role in the Organization of Microtubules. Int Rev Cytol 106: 227–293

Wei JH & Seemann J (2010) Unraveling the Golgi ribbon. Traffic 11: 1391–1400 doi:10.1111/j.1600-0854.2010.01114.x [PREPRINT]

Wu J & Akhmanova A (2017) Microtubule-Organizing Centers. Annu Rev Cell Dev Biol 33: annurev-cellbio-100616-060615

Wu J, de Heus C, Liu Q, Bouchet BP, Noordstra I, Jiang K, Hua S, Martin M, Yang C, Grigoriev I, et al (2016) Molecular Pathway of Microtubule Organization at the Golgi Apparatus. Dev Cell 39: 44–60

Yadav S, Puri S & Linstedt AD (2009a) A primary role for Golgi positioning in directed secretion, cell polarity, and wound healing. Mol Biol Cell 20: 1728–36

Yadav S, Puri S & Linstedt AD (2009b) A primary role for golgi positioning in directed secretion, cell polarity, and wound healing. Mol Biol Cell 20: 1728–1736

Yang C, Wu J, de Heus C, Grigoriev I, Liv N, Yao Y, Smal I, Meijering E, Klumperman J, Qi RZ, et al (2017) EB1 and EB3 regulate microtubule minus end organization and Golgi morphology. J Cell Biol 216: 3179–3198

